# Structural snapshots of V/A-ATPase reveal a new paradigm for rotary catalysis

**DOI:** 10.1101/2021.10.26.465817

**Authors:** J. Kishikawa, A. Nakanishi, A. Nakano, S. Saeki, A. Furuta, T. Kato, K. Mistuoka, K. Yokoyama

## Abstract

V/A-ATPase is a motor protein that shares a common rotary catalytic mechanism with F_o_F_1_ ATP synthase. When powered by ATP hydrolysis, the V_1_ moiety rotates the central rotor against the A_3_B_3_ hexamer, composed of three catalytic AB dimers adopting different conformations (AB_open_, AB_semi_, and AB_closed_). Here we have determined the atomic models of 18 catalytic intermediates of the V_1_ moiety of V/A-ATPase under different reaction conditions by single particle Cryo-EM, which revealed that the rotor does not rotate immediately after binding of ATP to the V_1_. Instead, three events proceed simultaneously with the 120° rotation of the shaft: hydrolysis of ATP in AB_semi_, zipper movement in AB_open_ by the binding ATP, and unzipper movement in AB_closed_ with release of both ADP and P*i*. This indicates the unidirectional rotation of V/A-ATPase by a ratchet-like mechanism owing to ATP hydrolysis in AB_semi_, rather than the power stroke model proposed previously for F_1_-ATPase.

## Main

The proton translocation ATPase/synthase family includes F-type enzymes found in eubacteria, mitochondria, and chloroplasts, and the V/A type enzymes found in archaea and some eubacteria^1–5^ (Fig. 1a). These ATPases produce the majority of cytosolic ATP from ADP and P*i* using energy derived from the transmembrane proton motive force generated by cellular respiration^6^. These ATPases share a common molecular architecture, consisting of a hydrophilic V_1_/F_1_ moiety responsible for ATP hydrolysis or synthesis, and a hydrophobic V_o_/F_o_ domain housing a proton translocation channel^7–9^. The chemical reaction (ATP hydrolysis/synthesis) in V_1_/F_1_ is tightly associated with proton movement through V_o_/F_o_ using a rotary catalytic mechanism, where both reactions are coupled by rotation of the central rotor complex relative to the surrounding stator apparatus, which includes the ATPase active hexamer^6, 10, 11^ (Fig. 1b).

**Fig. 1:**
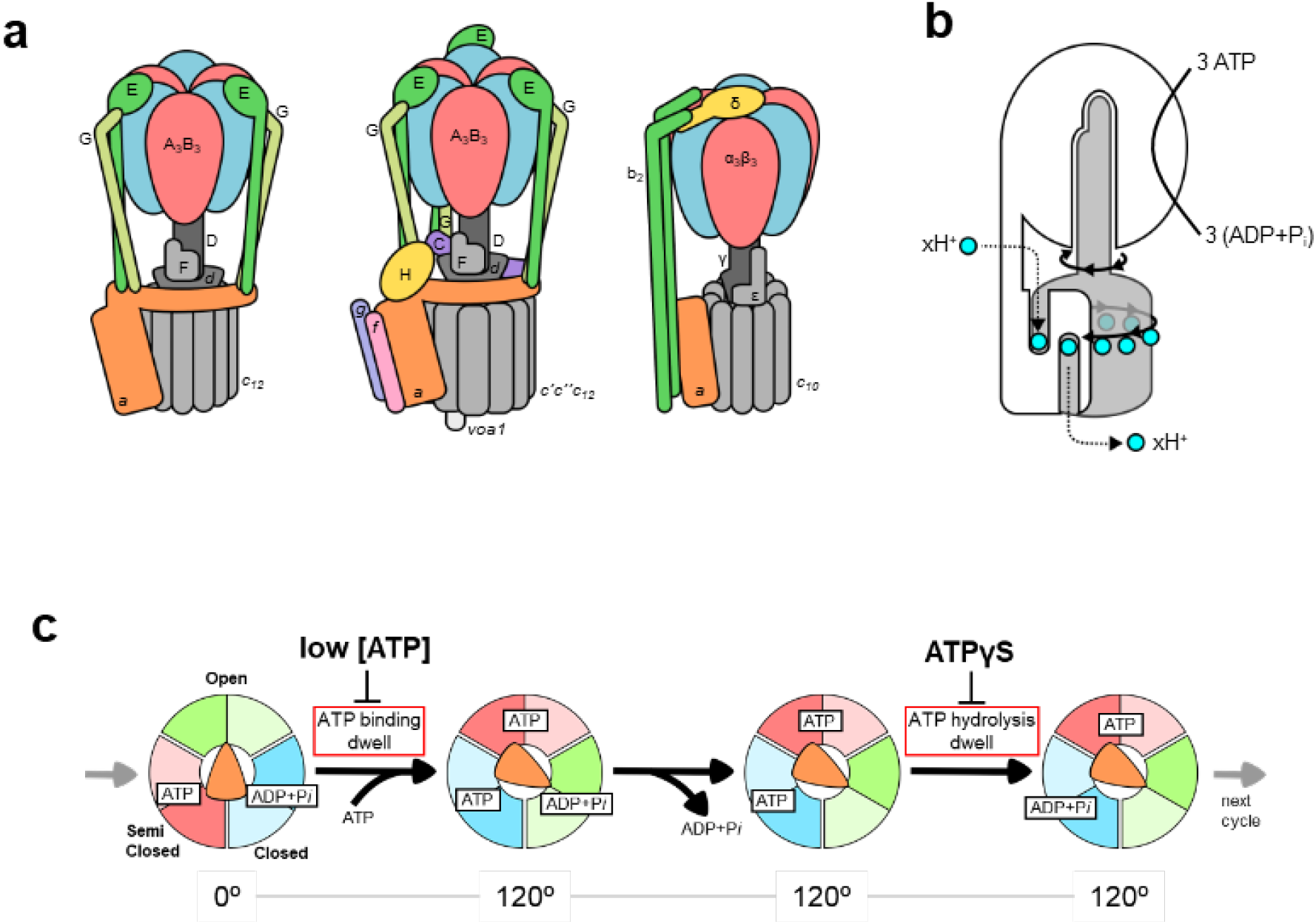
Schematic representation of rotary ATPases and the conventional rotary mechanism. **a**, Illustration of subunit composition of different types of rotary ATPases; prokaryotic V/A-ATPase (*left*), eukaryotic V-ATPase (*middle*), prokaryotic F-ATPase (*right*). The stators are represented in various colors and the rotors are represented in grey. **b**, A schematic model of the rotary catalytic mechanism of the V/A-ATPase. When powered by ATP, the central rotor composed of D_1_F_1_*d*_1_*c*_10_ (grey) rotates against a surrounding stator composed of A_3_B_3_E_2_G_2_*a*_1_ (white), coupled with proton translocation across the membrane. **c,** The conventional catalytic cycle of V/A-ATPase. At low ATP concentration, the ATP binding dwell time is increased. ATPγS also prolongs the ATP hydrolysis dwell.

The V/A-ATPase from the thermophilic bacterium, *Thermus thermophilus* (*Tth*) is one of the best-characterized ATP synthases^3, 12^. The overall architecture and subunit composition of V/A-ATPase is more similar to that of the eukaryotic V-ATPase, rather than F-type ATPase. However, the *Tth* V/A-ATPase has a simpler subunit structure than eukaryotic V-ATPase and shares the ATP synthase function of F-type ATPase^13^ (Fig. 1a). The V_1_ moiety of *Tth* V/A-ATPase (A_3_B_3_D_1_F_1_) is an ATP driven rotary motor where the central DF shaft rotates inside the hexameric A_3_B_3_ containing three catalytic sites, each composed of an AB dimer. The V_o_ domain (E_2_G_2_*d*_1_*a*_1_*c*_12_) is composed of stator parts including the *a* subunit and two EG peripheral stalks and the *d*_1_*c*_12_ rotor complex, which consists of a central rotor complex with the DF subunits of V_1_ ^14–16^. When ATP hydrolysis by A_3_B_3_ powers the DF shaft, the reverse rotation of the central rotor complex drives proton translocation in the membrane-embedded V_o_ domain (Fig. 1b).

According to the binding change mechanism of ATP synthesis^6^, the three catalytic sites in ATP synthases are in different conformations but interconvert sequentially between three different conformations as catalysis proceeds. Indeed, our previous structure demonstrated that the A_3_B_3_ hexamer in the V/A-ATPase adopts an asymmetrical structure composed of three different AB dimers, termed open (AB_open_), semi-closed (AB_semi_), and closed (AB_closed_)^16, 17^. The asymmetrical structure of the V_1_ implies that sequential interconversion between these three different AB dimers is synchronized with rotation of the central DF shaft. Experimental studies using specific rotational probes attached to DF, revealed that ATP-driven rotation of the central shaft was unidirectionally clockwise when viewed from the V_1_ side^10^. At low ATP concentrations where ATP binding is rate-limiting, rotation proceeds in steps of 120°, commensurate with the three catalytic sites of AB dimers ^18^. When using 40 nm gold beads with almost negligible viscous resistance, V_1_ also pauses every 120° even at a ATP concentration around *K*_m_ without a sign of substeps^19^. These single-molecule experiments on V_1_ suggest that the both catalytic events, ATP hydrolysis and product (ADP and P*i*) release occur at an individual ATP binding position, and imply the presence of chemo-mechanically stable catalytic intermediates (Fig. 1c and Supplementary Fig. 1).

However, single-molecule observation experiments only allow us to see the motion of the shaft to which the observation probe is bound, and do not tell us what events are occurring at each catalytic site. To elucidate the entire rotational mechanism of V/A-ATPase, we must determine the structures of catalytic intermediates of the rotary ATPase during rotation. There are many reaction intermediates of the enzyme during turnover, and this structural heterogeneity makes successful crystallization of a specific state very challenging.

Technological breakthroughs in single particle Cryo-EM, such as the development of direct electron detectors, and advances in image processing and automation^20, 21^, have triggered a revolution in structural biology, making this the technique of choice for large and dynamic complexes unsuitable for crystallization. In addition, by freezing Cryo-EM grids at different time points or under different reaction conditions, it is possible to trap intermediate states and thus build up a picture of the chemo-mechanical cycle of biological macromolecular complexes step by step. To date, there are few examples of studies that have successfully captured such details of a catalytic cycle at atomic resolution using Cryo-_EM_22,23.

Here we report several key, and thus far uncharacterized, intermediate states of V/A-ATPase, obtained under different reaction conditions. Comparison of these structures provides insight into a novel cooperativity between the three catalytic sites and demonstrates a rotary catalytic mechanism powered by ATP hydrolysis.

## Results

### Sample preparation for Cryo-EM structural analysis

We previously determined the Cryo-EM structures of the wild-type V/A-ATPase containing an ADP in the catalytic site of AB_closed_ ^16, 17^. The V/A-ATPase bound to the inhibitory ADP exhibits no ATPase activity until the ADP is removed^13, 16, 24^. Partial ADP removal from AB_closed_ is possible by dialysis against an EDTA-phosphate buffer, but it is difficult to obtain a homogenous nucleotide-free V/A-ATPase after such a treatment, due to the high binding affinity of the ADP to AB_closed_ (Supplementary Table 1). To obtain a homogeneous ATPase active enzyme, mutant V/A-ATPase (A/S232A, T235S) with reduced nucleotide-binding affinity was purified from *T. thermophilus* membranes^10^. The mutated V/A-ATPase exhibits higher *K*_m_ values for nucleotide in both the ATP hydrolysis and synthesis reactions than the wild-type enzyme, but the enzymatic and rotational properties are almost the same as those of the wild-type enzyme^24^. The mutated V/A-ATPase is fully activated for ATPase activity after EDTA/phosphate dialysis; no ADP or ATP was found in the enzyme by quantitative analysis of nucleotides (Supplementary Fig. 2 and Table 1). We incorporated the nucleotide-free V/A-ATPase into nanodiscs comprising the MSP1E3D1 scaffold protein and DMPC. The resulting V/A-ATPase obeys simple Michaelis-Menten kinetics and exhibits ATPase activity of 22 s^-1^ and the *K*_m_ of 394 µM ATP (Supplementary Fig. 2b).

The nucleotide-free V/A-ATPase (Nucfree) was used for cryo-grid preparation under different ATPase reaction conditions (Supplementary Fig. 1c). Results of the structural analysis of the protein under each set of reaction conditions are summarized in Supplementary Figs. 3a-d.

### The structures of V/A-ATPase without nucleotide (V_nucfree_)

The flow charts showing image acquisition and reconstitution of the 3D structure of V/A-ATPase without nucleotide are summarized in Supplementary Fig. 3a. We obtained structures of three rotational states of V/A-ATPase without nucleotide; state1 at 3.1 Å, state2 at 4.7 Å, and state 3 at 6.3 Å resolution, with the DF shaft positions differing by 120° in each case (Fig. 2). Using signal subtraction of the V_o_ domain, we achieved resolution of 3.0 and 4.1 Å for the V_1_ moiety including half of the EG stalk in state1 and 2, allowing us to build atomic models of the V_1_ domain of these states.

**Fig. 2:**
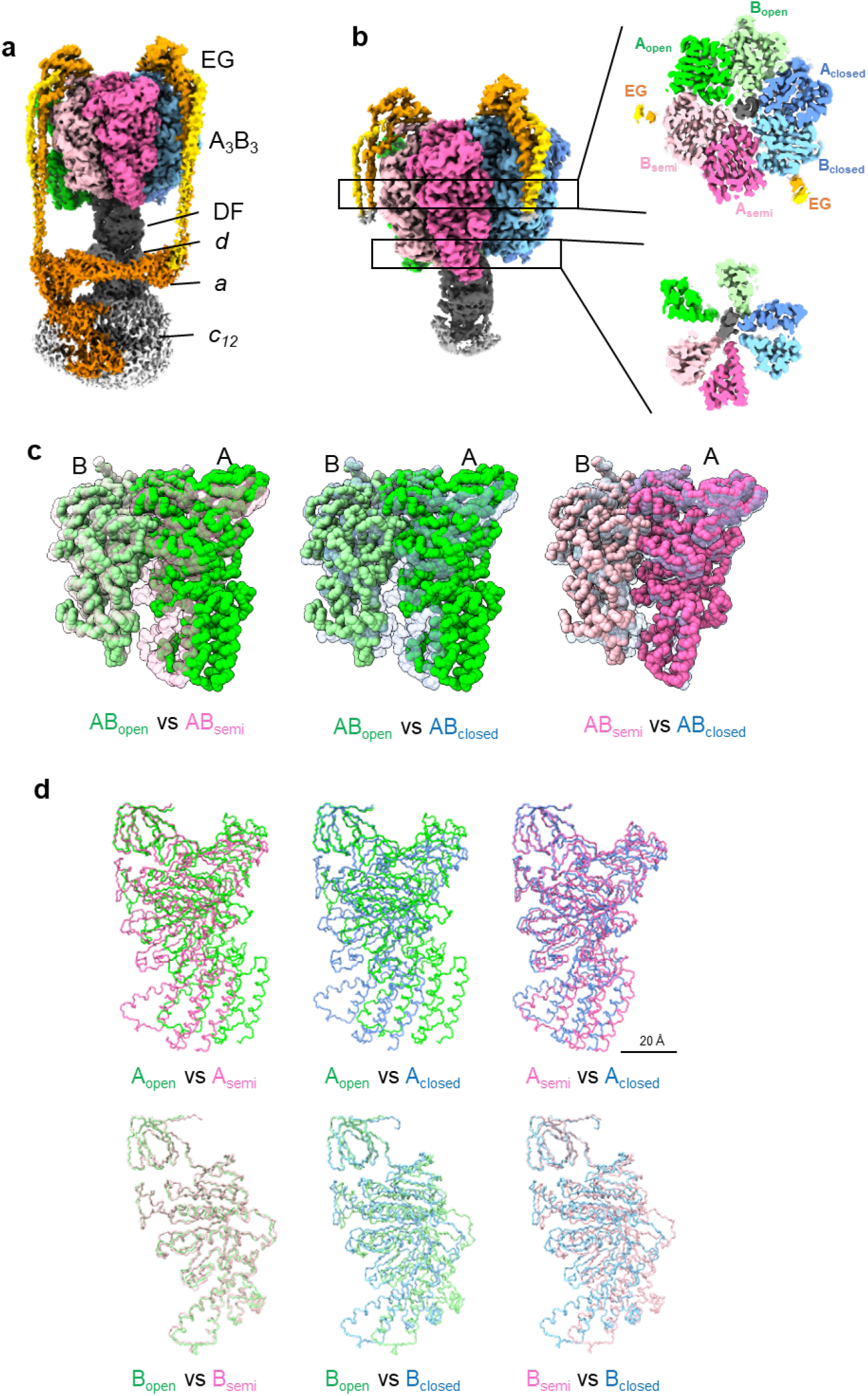
Cryo-EM density map and the atomic model for nucleotide free V/A-ATPase. **a**, Cryo-EM density map of whole V/A-ATPase of state1 in the absence of nucleotide (V_nucfree_ structure). **b**, Cryo-EM density map of V_1_EG of state1 without nucleotide (*left*). Cross sections of the nucleotide binding sites (*right upper*) and A_3_B_3_ C-terminal region (*right lower*) viewed from top. **c**, Comparison of the AB dimer structures in V_nucfree_. AB dimers are shown as space filling models and superimposed on the β barrel domain (A subunit 1-70 a.a.). *Left;* AB_open_ (solid) vs AB_semi_ (semi-transparent), *middle;* AB_open_ (solid) vs AB_closed_ (semi-transparent), and AB_semi_ (solid) vs AB_closed_ (semi-transparent). **d**, Comparison between each AB subunit in V_nucfree_. The subunits are shown as wire representation. A and B subunits are superimposed on the β barrel domain.

The three AB dimers in the V_1_ moiety adopted open (AB_open_), semi-closed (AB_semi_), and closed (AB_closed_) states, respectively (Fig. 2b and c). The tip of the C-terminal helix bundle (CHB) of A_open_ is in contact with the C-terminal helix of the D subunit, and the wide part of the CHB of B_open_ is in contact with the N-terminal helix of the D subunit, respectively (Supplementary Fig. 4 a-c). The AB_semi_ and AB_closed_ also interact with the coiled-coil of subunit D in specific regions of the CHB, respectively (Supplementary Fig. 4 d-g).

The differences in the structures of the three AB dimers when superimposed on the β barrel domains of both A and B subunits are the result of movement of the N-terminal bulge domain, the nucleotide binding domain (NB) of the A subunit and the CHBs of both the A and B subunits (Fig. 2d). When comparing the structure of AB_open_ and AB_semi_, both the NB and CHB of the A_semi_ are in closer proximity to B_semi_ than B_open_, resulting in a closed structure of AB_semi_ (Fig. 2c and d). The structure of B_open_ is very similar to B_semi_, as shown in Fig. 2d. In the AB_closed_, both the CHB and NB domains of A_closed_ are in closer proximity to B_closed_, and the CHB of B_closed_ moves to A_closed_, resulting in the more closed structure of AB_closed_ compared to AB_semi_ (Fig. 2c and d).

In the AB_closed_ and AB_semi_ dimers, densities for the catalytic side chains are well resolved, but no density corresponding to nucleotide was observed (Fig. 3a). Hereafter we refer to the structure as the V_nucfree_. The structure of V_nucfree_ is very similar to the previously reported ADP inhibited structure_16,17_. For state1, the *rmsd* value for the Cα chains of A_3_B_3_DF of the V_nucfree_ and ADP inhibited structures is 1.98 Å (Supplementary Fig. 5). In addition, the V_nucfree_ is also similar to the structures under saturated-ATP condition determined in this study, with the positions of the catalytic side chains almost identical in both cases (Supplementary Fig. 6). This indicates that the V_1_ moiety adopts the same conformation, including the arrangement of the DF shaft in the A_3_B_3_ and the geometry of the catalytic side chains, irrespective of the presence or absence of bound ATP.

**Fig. 3:**
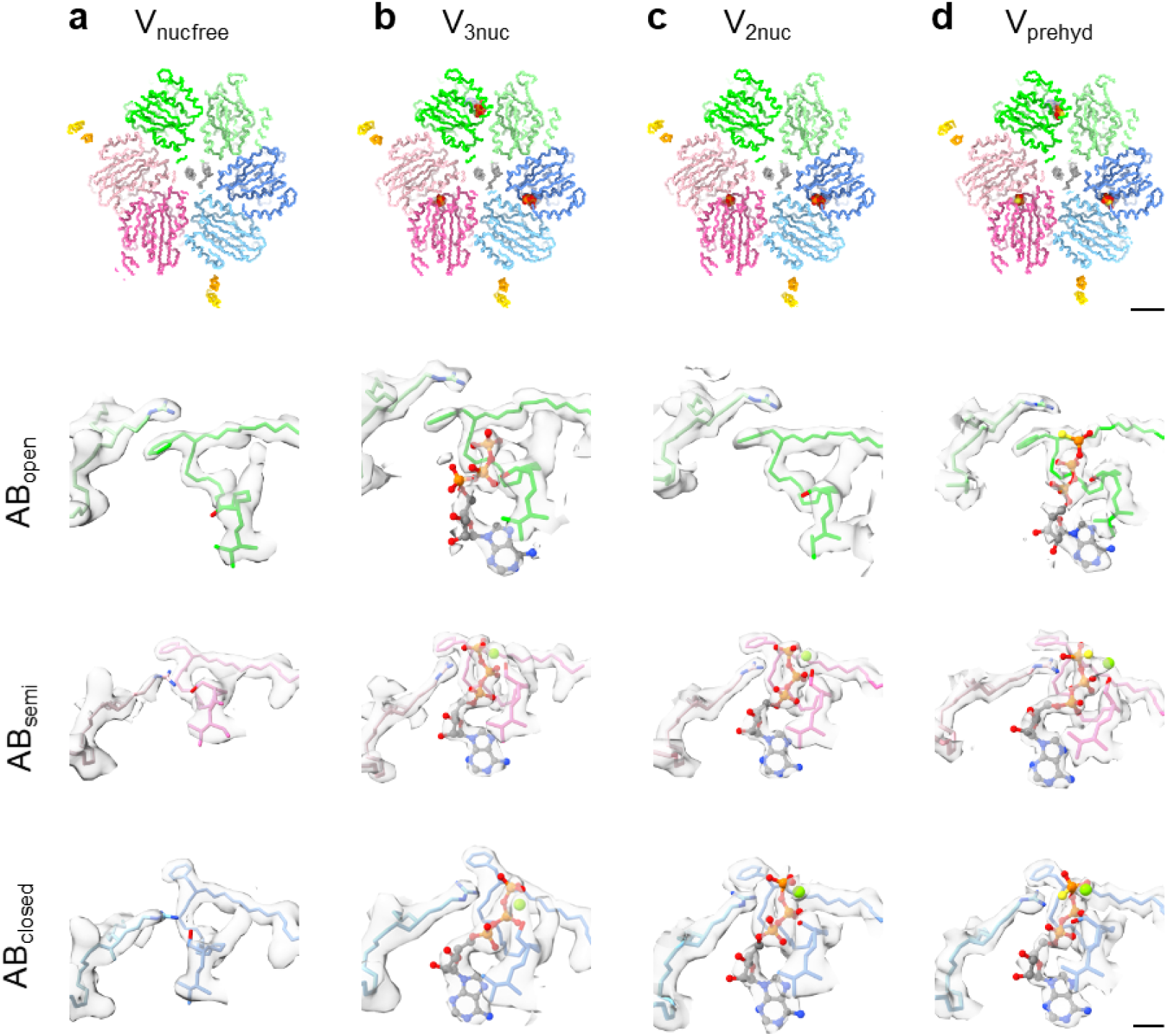
Structures of nucleotide binding sites obtained in each condition. *upper panels:* V_nucfree_ **(a)**, V_prehyd_ **(b)**, V_3nuc_ **(c)**, and V_2nuc_ **(d)** viewed from the cytosolic side. Scale bar is 20 Å. Magnified views of the three nucleotide binding sites (AB_open_, AB_semi_, and AB_closed_) in each structure are shown in the rows below. Cryo-EMmaps are represented as semi-translucent. Bound nucleotides and Mg ions are shown in ball-and-stick and sphere representation, respectively. Scale bar is 4 Å

In the density maps obtained for state1 of V_nucfree_, the CHB of the AB dimers were slightly blurred, likely due to structural heterogeneity. To classify the probable substates of state1, we performed focused 3D classification using a mask covering AB_open_ and B_semi_ (Supplementary Fig. 3a). We identified an atomic resolution structure of the original state1 from 39,902 particles at 3.1 Å resolution and another substate from 24,101 particles at 3.1 Å resolution. We termed the substates reconstructed from these major particle classes as state1-1 and state1-2, respectively. The atomic model initially constructed as state1 is identical to the atomic model of state1-1. The structures of the sub-states are very similar, with most differences due to the movement of the CHB of the A and B subunits. Therefore, we quantified the difference in the structure of the CHB observed when the structures were superimposed on the N-barrel domain (Supplementary Table 2 and 3). Substates were also obtained under other reaction conditions (see below) and the *rmsd* values shown in Supplementary Table 2 and 3 are used to discuss which subunits are responsible for the differences in structure of the substates obtained under different reaction conditions.

### Structures obtained at a saturating ATP concentration

Cryo-grids were prepared using a reaction mixture of Nucleotide-free V/A-ATPase, containing the regenerating system and ATP at a saturating concentration of 6 mM. The reaction mixture was incubated for 120 sec at 25 °C, and then loaded onto a holey grid, followed by flash freezing.

We determined three rotational states followed by focused refinement using a V_1_EG mask for each state (Supplementary Fig. 3b). In the density maps obtained for each state, the amino acid residues of the nucleotide binding sites in both AB_closed_ and AB_semi_ were well resolved, but the CHB domains of the AB dimers were blurred due to structural heterogeneity, as with the V_nucfree_. For state1, we identified an atomic resolution structure of state1-1 from 40,831 particles at 3.1 Å resolution and state1-2 from 28,801 particles at 3.2 Å resolution by further 3D classification without alignment (Supplementary Fig. 3b). The same classification analysis was performed for state2 and state3, yielding state 2-1 (3.0 Å resolution) and state 2-2 (3.4 Å resolution), and state 3-1 (3.0 Å resolution) and state 3-2 (3.4 Å resolution) respectively. In these structures, nucleotide densities have been identified in the three catalytic sites. Hereafter, we refer to the structures obtained at ATP saturating condition as V_3nuc_.

The structure of AB_open_ of V_3nuc_ state1-1 is almost identical to that of V_nucfree_ state1-1 (Supplementary Fig. 6). This is confirmed by the fact that the *rmsd* values in the CHB of A_open_ and B_open_ for V_3nuc_ state1-1 and V_nucfree_ are less than 1Å (Supplementary Table 2 and 3). The AB_open_ of state1-2 adopts a slightly more closed conformation compared to that of state1-1, which results from a movement of CHB of B_open_ towards the β-barrel domain (Fig. 4c). Nevertheless, the AB_open_ of V_3nuc_ with bound ATP retains the interaction with the DF shaft, indicating that ATP binding to the AB_open_ does not move the DF shaft.

**Fig. 4:**
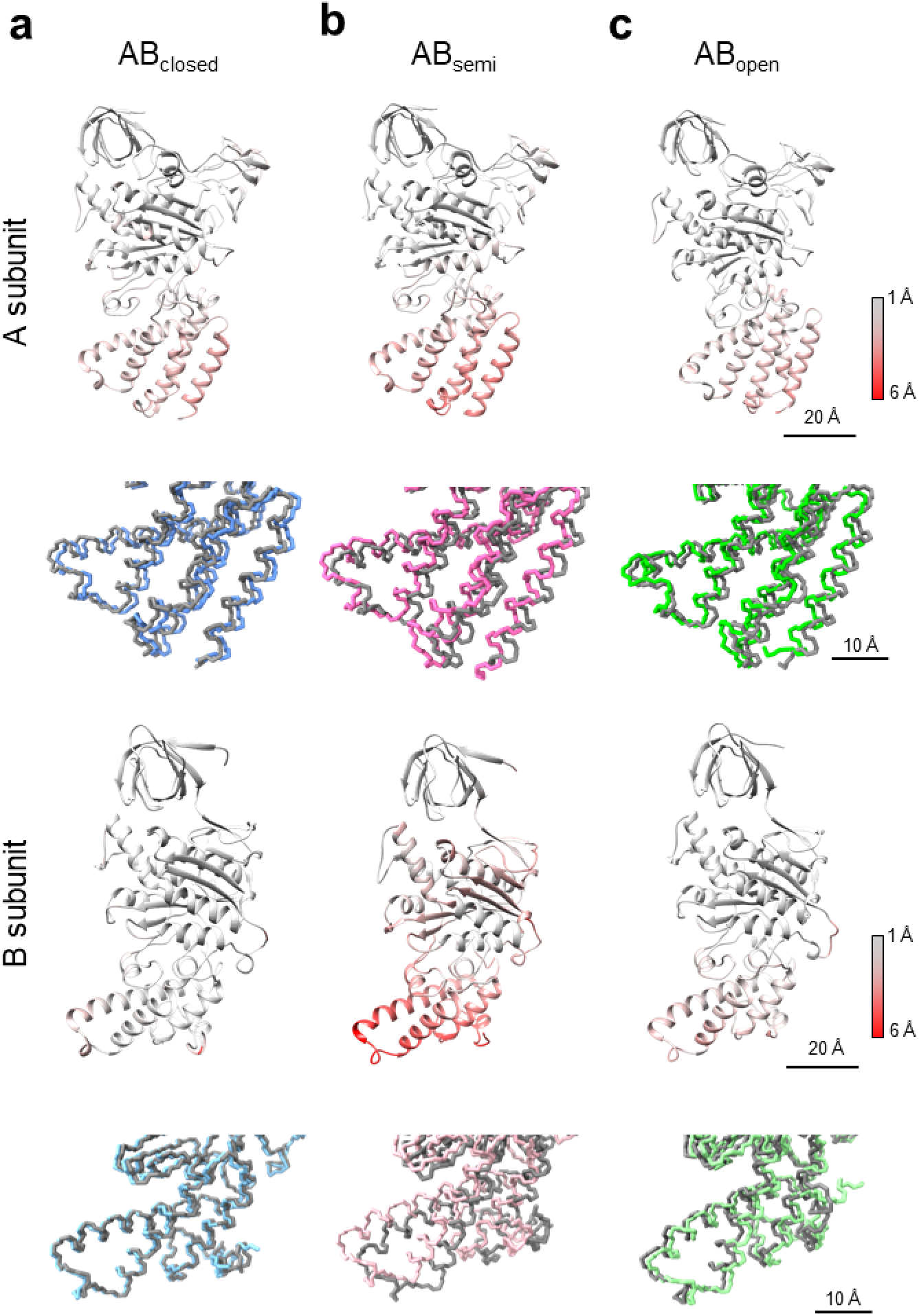
Comparison between state1-1 and 1-2 in each subunit of V_3nuc_. The subunits of state1-1 and 1-2 were superimposed on the β barrel domain (A: 1-70 a.a., B: 1-70 a.a.). Ribbon models are colored by the *rmsd* values calculated for the atoms of the main chain; gray (small changes) to red (large changes). Magnified views of the CHBs are represented in the lower panels as wire models. The models of state1-1 are represented in gray, and state1-2 are represented in the different colors. **a**, subunits in AB_closed_, **b**, subunits in AB_semi_, **c**, subunits in AB_open_.

The AB_semi_ in state1-2 has a more closed structure than that in state1-1 mainly due to the movement of CHB in B_semi_ (Fig. 4b, Supplementary Table 3). In summary, V_3nuc_ state1-2 has a more closed structure than state1-1 due to the movement of the CHB of both A_semi_ and B_semi_, but the slightly closed conformation of state1-2 is independent of ATP binding to the AB_open_.

### Structures of the catalytic sites at AB dimers of V_3nuc_

In both state1-1 and state1-2 structures obtained under ATP saturating conditions, a bound ATP molecule is clearly observed in the catalytic site of AB_open_ (Fig. 3b and Supplementary Fig. 7c). The catalytic sites in the AB_closed_ and AB_semi_ in both the state1-1 and state1-2 also contained density corresponding to an ATP molecule, and in these cases the associated magnesium ions were visible (Fig. 3b and Supplementary Fig. 7a-b). In the V_3nuc_ structure, we did not find density corresponding to nucleotides between the D and A subunits as reported in a previous paper_25_ (Supplementary Fig. 8).

In the catalytic site of AB_semi_ of V_3nuc_, the density of each nucleotide phosphate atom was easily identifiable (Supplementary Fig. 7b), indicating that the ATP molecule occupies the catalytic site in AB_semi_. The protein structure is sufficiently clear to also provide a detailed picture of the configuration of the catalytic side chains (Figs. 5a-c). The γ-phosphate of ATP and the magnesium ion are coordinated by the A/K234 and A/S235 residues on the P-loop, which contains the conserved nucleotide-binding motif_7,26_. The aromatic ring of A/F230, not conserved in F type ATPase, is oriented away from the triphosphate moiety, allowing access of the guanidium group to the arginine finger (Supplementary Fig. 9). Considering clear EM density for the γ-phosphate of the ATP bound in AB_semi_, hydrolysis of ATP is unlikely to proceed in AB_semi_.

**Fig. 5:**
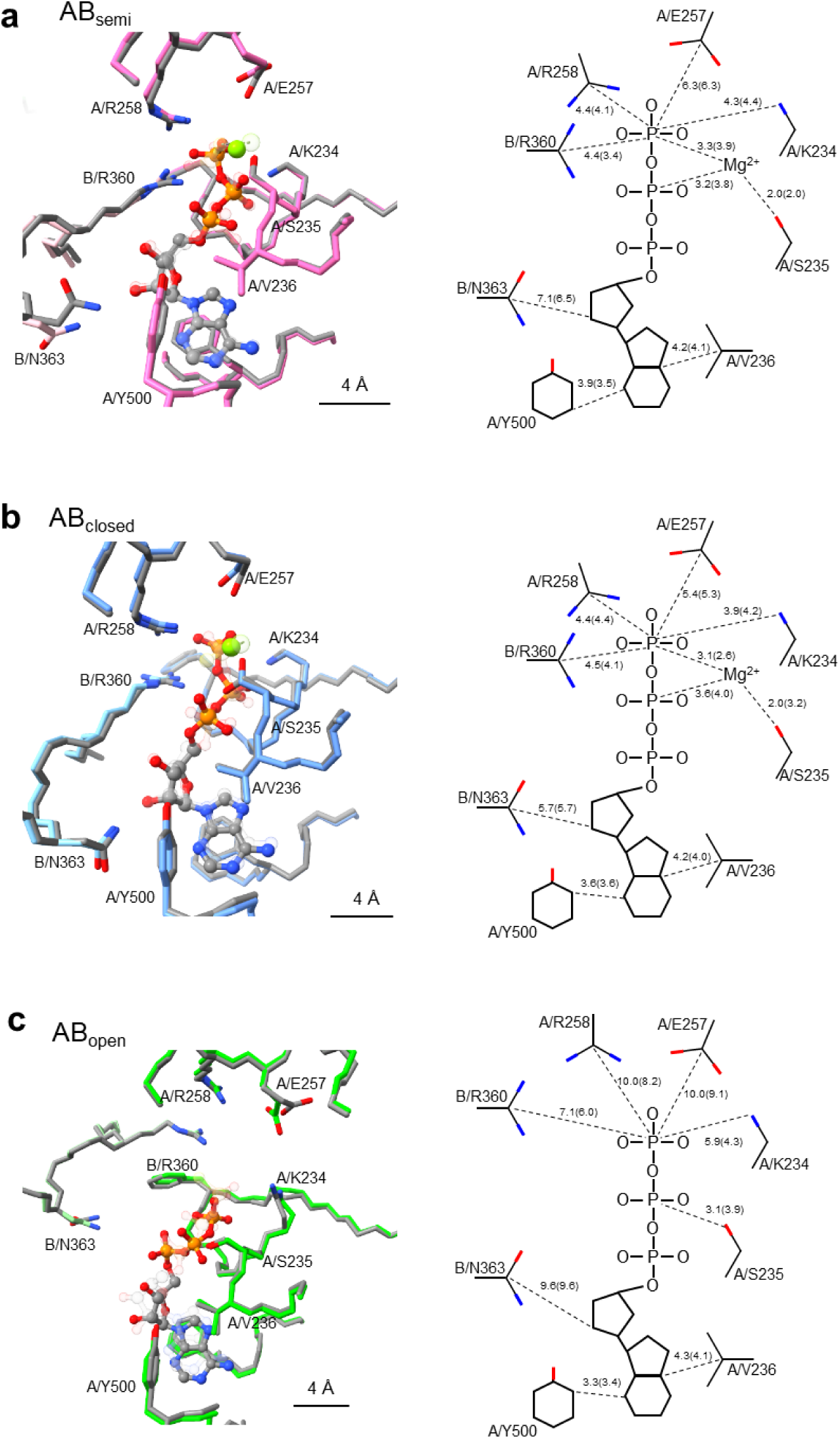
Coordination of nucleotides in the binding sites of V_3nuc_ and V_prehyd_. *Left panels;* Comparison of the three nucleotide binding sites (AB_open_ **(a)**, AB_semi_ **(b)**, and AB_closed_ **(c)**) of state1-1 of V_3nuc_ shown with colored (green, blue and pink) atoms and bonds, and main chain, and state1-1 of V_prehyd_ shown with grey atoms, bonds and main chain. *Right panels;* Schematic representations of the coordination of the ATP group in the three binding sites of V_3nuc_ and V_prehyd_ in parentheses. The distances between the atoms are shown in dotted lines. All distances are in Å.

The nucleotide-binding site of the AB_closed_ is shown in Fig. 5b. The geometry of ATP binding in AB_closed_ is very similar to that found in the AB_semi_, however, the carbonyl group of A/E257 is closer to the γ-phosphate by about 1 Å than AB_semi_. In state1-2 of V_3nuc_, the γ-phosphate of ATP bound to the AB_closed_ appears to be separated from β-phosphate when a relatively high density threshold is used (Supplementary Fig. 10). These findings strongly suggest that ATP bound to AB_closed_ is either already hydrolyzed or in the process of being hydrolyzed. The state of ATP in AB_closed_ is discussed further in Supplementary text.

In the nucleotide-binding site of the AB_open_ of V_3nuc_, the adenosine moiety of ATP is occluded as in the AB_semi_ and AB_closed_, with A/F415, A/Y500, and A/V236 forming the adenine binding pocket, however, the hydrogen bonding of the ribose moiety to the side chain of B/N363 is lost due to movement of the CHB of A_open_ (Fig. 5c). Unlike in the AB_closed_ and AB_semi_, the phenyl group of A/230F in the AB_open_ is closer to the tri-phosphate group of ATP due to torsion of the main chain, resulting in formation of a hydrophobic barrier between the catalytic side chains and the triphosphate moiety of ATP (Supplementary Fig. S9). Compared with AB_semi_, the side chains of catalytic residues A/E257, A/R258, and B/R360 of AB_open_ are much further away from the γ-phosphate of ATP (10.0, 10.0 and 7.1 Å, respectively. Consequently, the configuration of the catalytic residues in the nucleotide-binding site of AB_open_ is not appropriate for hydrolysis of the bound ATP. Instead, the bound ATP has the potential to zipper the AB interface via interaction with the surrounding catalytic residues, which ultimately results in the transition of AB_open_ to a more closed form via a typical zipper conformational change.

### Structures of V/A-ATPase waiting for ATP to bind

To determine the ATP-waiting structure of V/A-ATPase, we prepared a cryo-grid with a reaction mixture containing 4 µM enzyme, 50 µM ATP, and the ATP regeneration system, pre-incubated for 300 sec. We reconstructed three rotational states (state1, 2.7 Å, state2, 3.3 Å, and state3 3.6 Å resolution) from the single-particle images of the holo-complex and three rotational states of V_1_EG (state1, 2.8 Å, state2, 3.1 Å, and state3, 2.8 Å) by focused masked refinement. For state1, two substates (state1-1, 2.9 Å and state1-2, 3.0 Å) were separated by focused classification using AB_open_ and AB_semi_ masks (Supplementary Fig. 3c).

In both AB_semi_ and AB_closed_ of state1-1, apparent density of ATP-magnesium was observed, but the density of the γ-phosphate at the AB_closed_ is weaker than that at the AB_semi_ (Fig. 3c and Supplementary Figs. 7a and b). In contrast, density was not observed in the nucleotide-binding site of AB_open_. For state1-2, as in state1-1, nucleotides are present in both AB_semi_ and AB_closed_, while AB_open_ is empty. Hereafter we refer to the structure as V_2nuc_.

The overall structure and geometry of the catalytic residues of state1-1 of V_2nuc_ are largely identical to state1-1 of V_nucfree_ and V_3nuc_ (Supplementary Fig. 11). This structural similarity between V_2nuc_, V_nucfree_, and V_3nuc_ is confirmed by the low *rmsd* values when comparing the CHB of the A and B subunits of these structures (Supplementary Table 2 and 3). The similarity of the structures of these substates indicates that the structural polymorphism of V_1_ moiety is independent of the binding of ATP to AB dimers.

### Structures obtained at a saturating concentration of ATPγS (V_prehyd_)

The V_1_-ATPase from *T. thermophilus* is capable of hydrolyzing ATPγS, however, the turnover rate of ATPγS is much lower than that of ATP due to the decrease in hydrolysis rate^18^ (Supplementary Fig. 2d). Thus, pre-hydrolysis structures of V/A-ATPase can be obtained at 4 mM ATPγS. The cryo-grid was prepared by blotting of the reaction mixture comprising Nucleotide-free V/A-ATPase and 4 mM of ATPγS in the absence of the regenerating system in order to exclude any effect of regenerated ATP produced from hydrolyzed ATPγS. We reconstructed three rotational states from the acquired EM images using the CRYOARM300 (JEOL). After the focused masked refinement of the V_1_EG domain, we obtained atomic resolution structures of each state (state1, 2.7 Å, state2, 3.4 Å, and state3, 3.6 Å), respectively (Supplementary Fig. 3d). We refer to these structures as V_prehyd_. For state1, two sub-states, state1-1 and state1-2, were obtained at 2.7 and 2.9 Å resolution respectively, by focused classification using a mask with AB_open_ and AB_semi_ (Supplementary Fig. 3d). For the AB_open_ of V_prehyd_, bound ATPγS is clearly observed in the catalytic site, which has an almost identical structure to that of the ATP bound state of V_3nuc_ (Fig. 3b).

The nucleotide-binding sites of the AB_closed_ and AB_semi_ of V_prehyd_ are almost identical to those of V_3nuc_, respectively, as shown in Fig. 5. The γ-phosphate group of the bound ATPγS molecule in the AB_semi_ is well resolved as seen in for ATP in V_3nuc_ and V_2nuc_ (Figs. 3d and Supplementary Fig. 7b). In contrast, the density of γ-phosphate of ATPγS at the AB_closed_ is faint (Supplementary Fig. 7a), suggesting that the ATPγS in the AB_closed_ has already been hydrolyzed and the bound nucleotide in the AB_closed_ is ADP. This indicates that the AB_semi_ is in the pre-hydrolysis conformation, waiting for ATP hydrolysis.

## Discussion

### V/A-ATPase has a very similar structure irrespective of the nucleotide binding status

We have obtained catalytic intermediates of V_1_ moiety, V_nucfree_, V_3nuc_, V_2nuc_, and V_prehyd_, with different nucleotide occupancy. Despite the different nucleotide occupancy of these structures, their overall conformations are very similar. For instance, the *rmsd* of the Cα chains of A_3_B_3_ in V_nucfree_ and V_3nuc_ state1-1 is 1.98Å. In addition, the relative position of the central DF shaft within the asymmetric A_3_B_3_ is almost the same in the V_nucfree_ and V_3nuc_ structures. These findings demonstrate that the configuration between the DF shaft and individual AB dimers is independent of the state of nucleotide occupancy of each AB dimer. In other words, the structure of the V_1_ moiety adopts three rotational states, 1, 2, and 3, during continuous ATP hydrolysis, with the conformational changes in the A_3_B_3_ hexamer driven by ATP hydrolysis, being discrete rather than continuous.

### Chemo-mechanical cycle of the V/A-ATPase powered by ATP hydrolysis

The V_3nuc_ structure, obtained under ATP saturation conditions shows all three catalytic sites occupied by ATP or the products of hydrolysis (ADP + P*i*). Since the hydrolyzed P*i* is clearly visible in the AB_closed_ of V_3nuc_ (Supplementary Fig. 10), it is assumed that V_3nuc_ is the structure before dissociation of P*i* from the catalytic site in AB_closed_. We also obtained the

V_2nuc_ structure in which ATP and product(s) are bound to AB_semi_ and AB_closed_, respectively, but AB_open_ is empty. The V_2nuc_ is therefore assumed to be the structure of the protein awaiting ATP binding to AB_open_. When using ATPγS as a substrate, which has a very slow hydrolysis rate, the high-resolution atomic structure of the V_1_ moiety allowed visualization of ATPγS molecules bound to the catalytic sites of AB_open_ and AB_semi_, as well as identification of the hydrolyzed ATPγS at the AB_closed_. The V_prehyd_ reveals both that the AB_closed_ adopts the post-hydrolysis state where the product of phosphate (P*i*) is dissociated, and that the AB_semi_ is awaiting ATP hydrolysis.

The structures provide important insights into the chemo-mechanical cycle of V/A-ATPase. The V/A-ATPase undergoes a unidirectional conformational change from state1 to state2 to state3 when powered by ATP. Thus, V_3nuc_ of state1, in which three catalytic sites are already occupied by nucleotides, should change to state2 of V_2nuc_, following ATP hydrolysis at AB_semi_, and the subsequent or simultaneously dissociation of ADP and P*i* by the discrete structural transition of AB_closed_ to AB_open_ (Fig. 6). This demonstrate that rotation of the rotary ATPase proceeds via the tri-site model with the protein progressing through a two nucleotide bound state and a three nucleotide bound state, settling the long-standing debate on whether the bi-site model or tri-site model is appropriate for rotary ATPases_5,6,27-30._

**Fig. 6:**
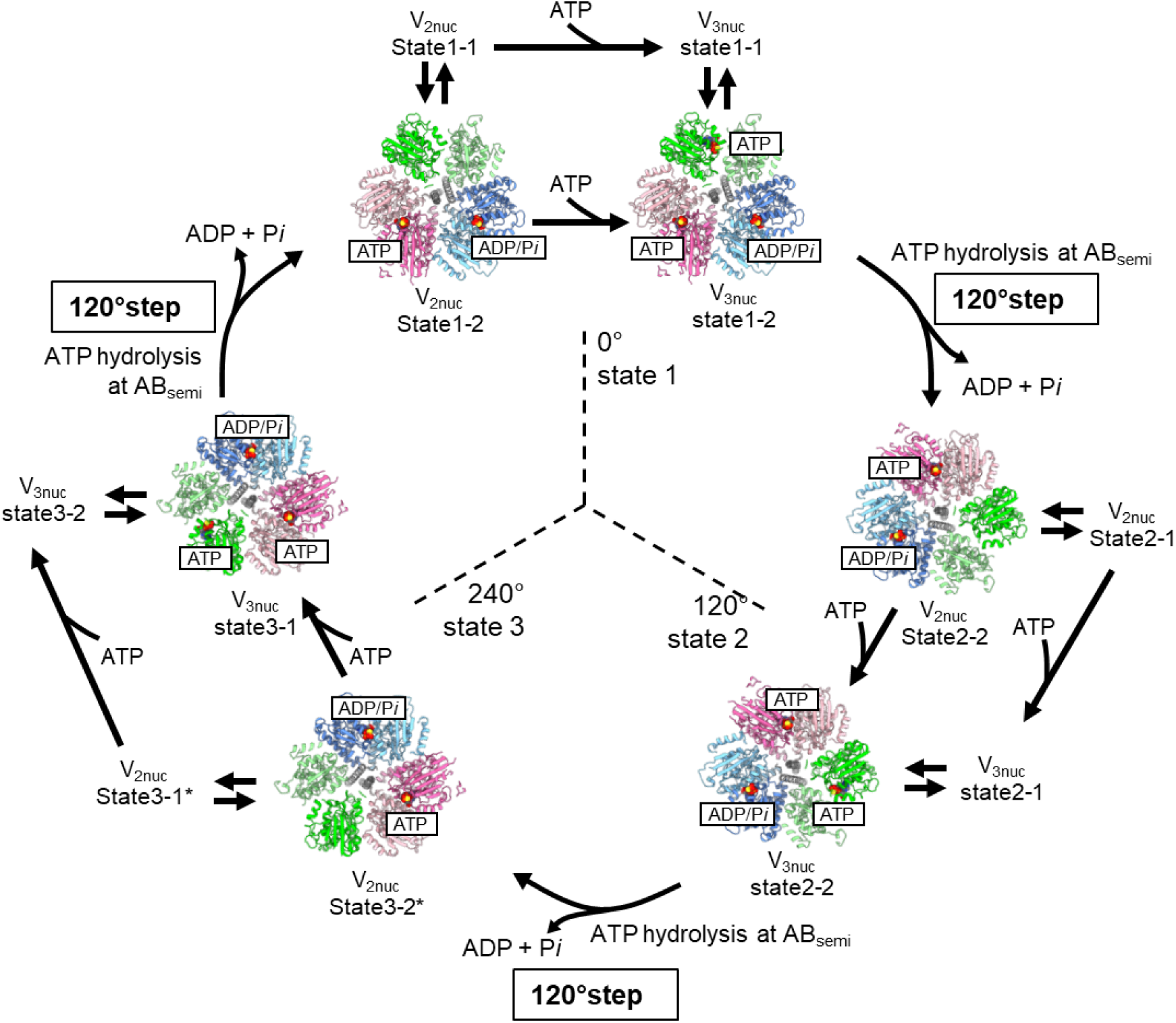
Chemo-Mechanical cycle of V/A-ATPase driven by ATP hydrolysis. The structures of V/A-ATPase viewed from the cytosolic side are shown as ribbon models. The coiled coil of the DF subunits is shown in grey. The bound ATP molecules are highlighted in sphere representations. State1-1 and 1-2 of V_2nuc_ are in equilibrium and are fluctuating. These structures transit to state 1-1 and 1-2 of V_3nuc_ by ATP binding to AB_open_, without a 120° rotation step of the DF rotor. V_3nuc_ in state1-1 and state1-2 are also in equilibrium. ATP hydrolysis at AB_semi_ and zipper motion at AB_open_ occur simultaneously. This triggers the transition of V_3nuc_ in state1-2 to V_2nuc_ state2-2 together with the 120°rotation step and simultaneous release of ADP and P*i*. State2 of V_2nuc_ returns to state1 via state3 of V_2nuc_ by the same process. Asterisks indicate the structures which were not identified in this study.

Based on previous single molecular observation experiments for both F_1_- and V_1_-ATPase, ATP binding onto the enzyme directly triggers the first 120° rotation step of the DF shaft_18,19,30_ (Supplementary Fig. 1a). In light of our findings presented here, this scheme needs to be redrawn; the rotor does not immediately travel 120° as a result of ATP binding to enzyme.

The next catalytic event after ATP binding is ATP hydrolysis in AB_semi_. Each conformational change, from AB_open_ to AB_semi_, AB_semi_ to AB_closed_, and AB_closed_ to AB_open_ occurs simultaneously, with the rotation of the shaft, and with the hydrolysis of ATP in the AB_semi_ and release of products (ADP and P*i*) from the AB_closed_ (Fig. 6 and Supplementary Movie 1). This is in marked contrast to the classical rotary model, where catalytic events occur in sequence at the three catalytic sites, until now the broadly accepted mechanism of action of the F_1_-ATPase^6, 27, 31, 32^.

In the V_2nuc_ and V_3nuc_, two sub-states, state1-1 and state1-2 were identified. These substates were also identified in V_nucfree_, thus the conformational dynamics of the V_1_ moiety are independent of ATP binding. In other words, state1-1 and state1-2 are in thermally equilibrium state, irrespective of nucleotide occupancy in each catalytic site. Both AB_semi_ and AB_open_ in state1-2 adopt more closed structures than those in state1-1, suggesting that state1-2 of V_3nuc_ is likely an intermediate structure just prior to the 120° rotation step of the DF shaft. Compared to state1-2 of V_3nuc_, state1-2 of V_prehyd_ exhibits slightly more closed structures of AB_open_ and AB_semi_ (Supplementary Fig. 13), likely to be associated with the progress of the catalytic reaction in AB_closed_, i.e., the dissociation of the phosphate. In this respect, state1-2 of V_prehyd_ may be another reaction intermediate structure in which the P*i* in the AB_closed_ is released prior to the 120° step (Supplementary Fig. 14).

### Rotation Mechanism of the V/A-ATPase

Based on the catalytic intermediates of the V_1_ moiety of V/A-ATPases obtained under four different reaction conditions, we propose a model for the ATP-driven rotation mechanism of V/A-ATPases.

When ATP binds to V_nucfree_, which is in a stable initial state (ground state), the enzyme transits into the steady state for ATP hydrolysis (Fig. 7, *upper row*). The V_3nuc_ structure is formed by binding of ATP to the AB_open_ of V_2nuc_ but the binding of ATP itself does not cause structural transitions between AB dimers associated with the 120° step of the DF shaft.

**Fig. 7:**
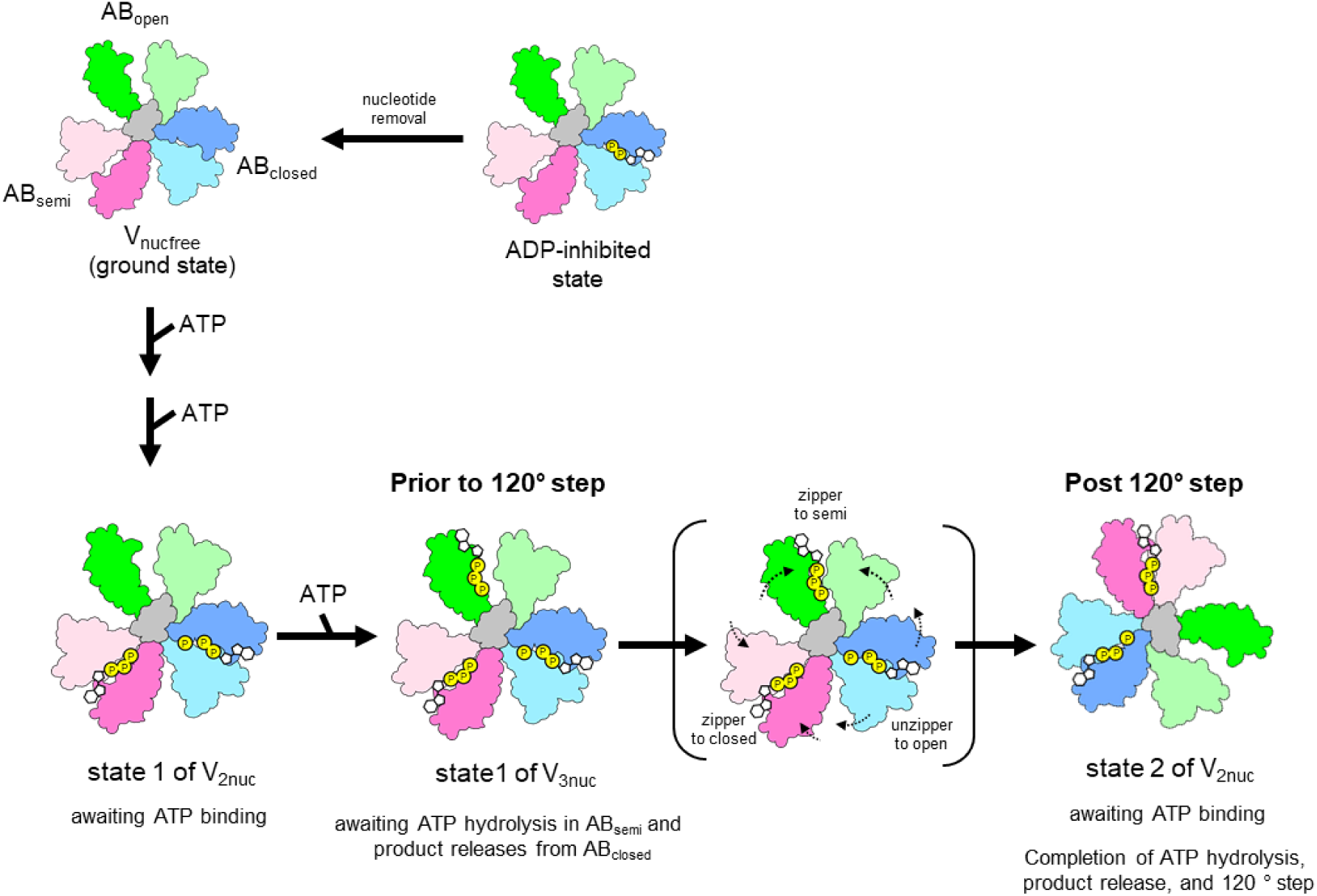
The rotary mechanism of V/A-ATPase powered by ATP hydrolysis. The schematic models of AB_open_, AB_semi_, and AB_closed_ are shown in green, pink, and blue, respectively. The coiled coil region of the D subunit in contact with A_3_B_3_ is shown in grey. In the ADP inhibited state, the entrapped ADP in AB_closed_ hampers structural transition of AB_open_ to AB_semi_ by binding of ATP to AB_open_. The V_nucfree_ in ground state is activated by the binding of ATP to the catalytic sites. In V_2nuc_ awaiting ATP binding, binding of ATP to AB_open_ does not induce the 120° rotation step. In V_3nuc_, both zipper motion of AB_open_ and ATP hydrolysis in AB_semi_ induce unzipper motion of AB_closed_ accompanying with the release of ADP and P*i*. The catalytic events in the three AB dimers occurs simultaneously with the 120° step of the DF shaft, resulting in structural transition of state1 of V_3nuc_ to state2 of V_2nuc_.

In the V_3nuc_ structure, where nucleotides are bound to all three AB dimers, three distinct but associated catalytic events occur at the three AB dimers simultaneously and these events are coupled to the first 120° rotation step of the DF shaft. One of the driving events for this transition is the conformational change from the ATP-bound AB_open_ to the more closed AB_semi_, which can be explained by a zipper motion of AB_open_ occurring upon ATP binding. From our structures, a comparison between AB_open_ and AB_semi_ implies that new hydrogen bonds form between the triphosphate moiety of ATP and the surrounding side chain groups of B/R360, A/258R, A/257E, and A/K234. (Fig. 5).

The V_prehyd_ structures indicate that the ATP bound to AB_semi_ is awaiting hydrolysis. The conformational change from AB_semi_ to AB_closed_ should occur spontaneously because it involves ATP hydrolysis, an exergonic reaction. In contrast, ADP bound to AB_closed_ hampers the unzipper motion in AB_closed_, thereby preventing the overall structural transition of the V_1_ moiety. This is supported by the fact that V/A-ATPase adopts the ADP inhibited state in which ADP is entrapped in AB_closed_ (Fig. 7, upper line). The enzyme in the ADP-inhibited state does not show ATP hydrolysis activity even at saturated ATP concentration_13, 16_. In summary, the ATP-driven unidirectional rotation of V/A-ATPase proceeds by a discrete structural transition between the three rotational states, i.e., the potential barrier to the structural transition of AB_closed_ to AB_open_, accompanied by release of ADP and P*i*, is overcome by both a zipper motion of AB_open_ by the bound ATP and ATP hydrolysis in AB_semi_. Since the ATP hydrolysis reaction is a heat dissipation process, the structural transition of AB_semi_ to AB_closed_ associated with the ATP hydrolysis occurs spontaneously and irreversibly, resulting in a unidirectionality of the 120° steps of rotor. In other words, our model explains the unidirectional rotation by a ratchet-like mechanism driven by ATP hydrolysis, rather than the power stroke model proposed previously for F_1_-ATPase^5, 33^.

V/A-ATPase and F_o_F_1_ are molecular machines based on the same construction principle, and thus are likely to share the same rotary mechanism. Importantly, for the thermophilic F_1_, the first 120° rotation step also includes ∼80° and ∼40° substeps, suggesting the existence of at least one additional catalytic intermediate of F1^5, 27, 34^. A recent structural study of thermophilic F_1_-ATPase indicated a possible intermediate structure responsible for the substeps^32^. In the V/A-ATPase, any intermediate structure containing phosphate after ADP or P*i* release is likely to be an unstable state, and therefore studies on the ATP driven rotation of V/A-ATPase have failed to reveal the presence of any substep^19^.

## Online Methods

### Preparation of *Tth* V/A-ATPase for Biochemical assay and Cryo-EM imaging

The *Tth* V/A-ATPase containing His3 tags on the C-terminus of each c subunit and the TSSA mutation (S232A and T235S) on the A subunit was isolated from *Thermus thermophilus* membranes as previously described^24^ with the following modifications. The enzyme, solubilized from the membranes with 10 % Triton X-100 was purified by Ni^2+^-NTA affinity with 0.03 % dodecyl-β-D-maltoside (DDM). For bound nucleotide removal, the eluted fractions containing *Tth* V/A-ATPase were dialyzed against 200 mM Sodium phosphate, pH 8.0, 10 mM EDTA, and 0.03% DDM over night at 25°C with three buffer changes, followed by dialysis against 20 mM Tris-Cl, pH 8.0, 1 mM EDTA, and 0.03% DDM (TE buffer) prior to anion exchange chromatography using a 6 ml Resource Q column (GE healthcare). The *Tth* V/A-ATPase was eluted by a linear NaCl gradient using a TE buffer (0-500 mM NaCl, 0.03% DDM). The eluted fractions containing *holo*-*Tth* V/A-ATPase were concentrated to ∼ 10 mg/ml using Amicon 100 k molecular weight cut-off fulters (Millipore). For nanodisc incorporation, the 1,2-Dimyristoyl-sn-glycero-3-phosphorylcholine (DMPC, Avanti) was used to form lipid bilayers in reconstruction as previously described^16^. Purified *Tth* V/A-ATPase solubilized in 0.03 % n-Dodecyl-β-D-maltoside (DDM) was mixed with the lipid stock and membrane scaffold protein MSP1E3D1 (Sigma) at a specific molar ratio V_o_V_1_: MSP: DMPC lipid = 1: 4: 520 and incubated on ice for 0.5 h. Then, 200 μL of Bio Beads SM-2 equilibrated with a wash buffer (20 mM Tris-HCl, pH8.0, 150 mM NaCl) was added to the 500 μL mixture. After 2 hours incubation at 4°C with gentle stirring, an additional 300 μL of Bio Beads was added and the mixture incubated overnight at 4°C to form the nanodiscs. The supernatant of the mixture containing nanodisc-*Tth* V/A-ATPase (nd-V/A-ATPase) was loaded onto the Superdex 200 Increase 10/300 column equilibrated with wash buffer. The peak fractions were collected, analyzed by SDS-PAGE and concentrated to ∼4 mg/mL. The prepared *nd*-V/A-ATPase was immediately used for biochemical assay or cryo-grid preparation, since *nd*-V/A-ATPase aggregates within a few days.

### Biochemical assay

The quantitative analysis of bound nucleotides of *Tth* V/A-ATPase was carried out using anion exchange high performance liquid chromatography^13^. Bound nucleotides were released from the enzyme by addition of 5 μl of 60% perchloric acid to 50 μl of the enzyme solution. Thereafter, the mixture was incubated on ice for 10 min. Then, 5 μl of 5 M K_2_CO_3_ solution was added and the mixture incubated on ice for 10 min. The resulting pellet was removed by centrifugation at 4°C. The supernatant was applied to a Cosmopak-200 column equilibrated with 0.1 M sodium-phosphate buffer (pH 7.0). The column was eluted isocratically with the same buffer at a flow rate of 0.8 ml/min. The nucleotide was monitored at 258 nm. The peak area was determined by automatic integration.

ATPase activity was measured at 25°C with an enzyme-coupled ATP-regenerating system, as described previously^13^. The reaction mixture contained 50 mM Tris-HCl (pH 8.0), 100 mM KCl, different concentrations of ATP-Mg, 2.5 mM phosphoenolpyruvate (PEP), 50 μg/ml pyruvate kinase (PK), 50 μg/ml lactate dehydrogenase, and 0.2 mM NADH in a final volume of 2 ml. The reaction was started by addition of 20 pmol *nd* V/A-ATPase to 2 ml of the assay mixture, and the rate of ATP hydrolysis monitored as the rate of oxidation of NADH determined by the absorbance decrease at 340 nm.

### Cryo-EM imaging of *Tth* V/A-ATPase

Sample vitrification was performed using a semi-automated vitrification device (Vitrobot, FEI). For *nd*-V/A-ATPase that underwent nucleotide removal, hereafter referred as nucfree *nd*-V/A-ATPase, 2.4 μl of sample solution at a concentration of 3 mg/ml (2 μM) was applied to glow discharged Quantifoil R1.2/1.3 molybdenum grid discharged by Ion Bombarder (Vacuum Device) for 1 min. The grid was then automatically blotted once from both sides with filter paper for 6 s blot time. The grid was then plunged into a liquid ethane with no delay time.

The reaction basal buffer (RB buffer) containing 50 mM Tris-Cl, pH 8.0, 100 mM KCl, and 2 mM MgCl_2_ was used for different reaction conditions. For saturated ATP or ATP waiting condition, 4 µM of nucfree *nd*-V/A-ATPase was mixed with same volume of x2 RB buffer containing 10 mM PEP, 200 μg/ml of PK, 12 mM or 100 µM of ATP-Mg. Then the mixtures were incubated for 120 sec or 300 sec at 25 °C, followed by blotting and vitrification, respectively. For the ATPγS saturated condition, 4 µM of nucfree *nd*-V/A-ATPase was mixed with the same volume of x2 RB buffer containing 8 mM ATPγS-Mg, then incubated for 300 sec at 25 °C, followed by the blotting and vitrification.

With the exception of the saturated ATPγS condition, cryo-EM imaging was performed with a Titan Krios (FEI/Thermo Fisher) operating at 300kV acceleration voltage and equipped with a direct K3 (Gatan) electron detector in electron counting mode (CDS). Data collection was carried out using SerialEM software^35^ at a calibrated magnification of 0.88 Å pixel^-1^ (x 81,000) and total dose of 50.0 e^−^ Å^-2^ (or 1.0 e^−^Å^-2^ per frame) (where e^−^ specifies electrons) with total 5 s exposure time. The defocus range was -0.8 to -2.0 μm. The data were collected as 50 movie frames.

For the saturated ATPγS condition, Cryo-EM movie collection was performed with a CRYOARM 300 (JEOL) operating at 300 keV accelerating voltage and equipped with a K3 (Gatan) direct electron detector, in electron counting mode (CDS) using the data collection software serialEM. The pixel size was 1.1 Å/pix (x60,000) and a total dose of 50.0 e^-^ Å^-2^ (1.0 e^-^ Å^-2^ per frame) with a total 3.0 s exposure time (50 frames) with a defocus range of -1.0 to -3.5 μm.

### Image processing

Image processing steps for each reaction condition are summarized in Figure S3 A-D. Image analysis software, Relion 3.1 and Cryosparc 3.2, were used^36, 37^. CTFFIND 4.1 and MotionCor2 were used for CTF estimation and movie correction in Relion^38, 39^. Topaz software was used for machine-learning based particle picking^40^. We started with 15,317 movies for the nucleotide free enzyme (nucfree *nd*-V/A-ATPase), 13,164 movies for saturated ATP condition, 15,711 movies for saturated ATP condition, and 17,522 movies for ATPγS condition. The software used in the steps is indicated in the figure. Autopicking based on template matching or based on Topaz machine-learning resulted in 4,354,341 particles for the nucfree *nd*-V/A-ATPase, 2,300,834 particles for the ATP saturated condition, 1,671,397 particles for the ATP waiting condition, and 4,677.284 particles for the ATPγS waiting condition. Particles were extracted at 5x the physical pixel size from the movie-corrected micrographs and selected using 2D or 3D classification (nucfree nd-V/A-ATPase; 132,904 particles, saturated ATP; 188,673 particles, ATP waiting; 186,928 particles, ATPγS; 197,960 particles). The selected particles were extracted at full pixel size and subjected to 3D auto-refinement refollowed by CTF refinement by Bayesian polishing. Another round of 3D auto-refine, CTF refinement, and a final round of masked auto-refinement gave *holo*-V/A-ATPase maps at between 2.7 ∼ 6.3 Å resolution. The membrane domain was visible but not particularly clear compared to the hydrophilic V_1_ domain in a *holo-*enzyme map. This seemed to be due to the structural flexibility between the membrane domain and V_1_ in the *holo*-enzyme. Focused refinement with signal subtraction targeting the V_1_EG region improved the map quality of the V_1_EG region (Figure S3A-D). The refinements provided the density maps for V_1_EG under each condition at 2.8 - 4.1 Å resolution. After the focused refinement, masked classification on A_open_ and B_semi_ subunits was carried out to classify the conformational differences. Resolution was based on the gold standard Fourier shell correlation = 0.142 criterion.

### Model building and refinement

To generate the atomic model for the V_1_EG region of V/A-ATPase, the individual subunits of the V_1_EG model from the previous structure of V/A-ATPase (PDBID: 6QUM) were fitted into the density map as rigid bodies^25^ with particular focus on the N terminal region of EG stalk (E; 1-77 aa., G; 2-33 aa.). The rough initial model was refined against the map with Phenix suite phenix.real_space_refine program^41^. The initial model was extensively manually corrected residue by residue in COOT^41^ in terms of side-chain conformations. Peripheral stalks were removed due to low resolution in this region. The corrected model was again refined by the phenix.real_space_refine program with secondary structure and Ramachandran restraints, then the resulting model was manually checked by COOT. This iterative process was performed for several rounds to correct remaining errors until the model was in good agreement with geometry, as reflected by the MolProbity score of 1.08∼1.74 and EMRinger score of 1.59 ∼3.94^42, 43^. For model validation against over-fitting, the built models were used for calculation of FSC curves against both half maps, and compared with the FSC of the final model against the final density map used for model building by phenix.refine program. The statistics of the obtained maps and the atomic model were summarized in Table S4. RMSD values between the atomic models were calculated using UCSF chimera^44^. All the figures were rendered using UCSF chimeraX^45^.

## Acknowledgements

We are grateful to all the members of the Yokoyama Lab for their continuous support and technical assistance. Our research was supported by Grant-in-Aid for Scientific Research (JSPS KAKENHI) Grant Number 20H03231 to K.Y., 20K06514 to J.K., and Grant-in-Aid for JSPS Fellows Grant Number 20J00162 to A.Nakanishi, and Takeda Science foundation to K.Y. Our research was also supported by Platform Project for Supporting Drug Discovery and Life Science Research (Basis for Supporting Innovative Drug Discovery and Life Science Research (BINDS)) from AMED under Grant Number JP17am0101001 (support number 1312), Grants-in-Aid from “Nanotechnology Platform” of the Ministry of Education, Culture, Sports, Science and Technology (MEXT) to K.M. (Project Number. 12024046), and the Research Program for Next Generation Young Scientists of "Five-star Alliance" in "NJRC Mater. & Dev." under Grant Number 20215008 to A. Nakano.

## Author contributions

KY, JK, A.Nakanishi and A.Nakano designed, performed and analyzed the experiments. JK, A.Nakanishi, KY, A.Nakano, AF, and SS analyzed the data and contributed to the preparation of the samples. TK and KM provided technical support and conceptual advice. KY designed and supervised the experiments and wrote the manuscript. All authors discussed the results and commented on the manuscript.

## Declaration of interests

The authors declare no conflicts of interest associated with this manuscript.

## Notes

### Competing Interest Statement

The authors have declared no competing interest.

