## Supplementary information for "Structural snapshots of V/A-ATPase reveal a new paradigm for rotary catalysis"

#### Supplementary text

##### Catalytic state of nucleotide bound to $AB_{closed}$

In the $V_{3nuc}$ structure, the EM density of the γ-phosphate of ATP in $AB_{closed}$ is clearer than that in the $V_{prehyd}$ structure (Figs 3 and Supplementary Figure 7). This difference is due to the length of the dwell time of the nucleotide in the catalytic site in the two different structures. The weak density of the γ-phosphate in the $AB_{closed}$ of $V_{prehyd}$ is likely due to the release of the cleaved phosphate during the slower hydrolysis reaction that occurs with ATPγS as a substrate. Since the ATP-binding dwell time is negligible under ATP-saturated conditions, $V_{3nuc}$ structures likely reflect the structures during and/or just after ATP hydrolysis in $AB_{closed}$. In fact, γ-phosphate of ATP bound to the $AB_{closed}$ of state1-2 of $V_{3nuc}$ appears to be separated from β-phosphate (Supplementary Figure 10). In addition, the density of the γ-phosphate of ATP bound to $AB_{closed}$ in $V_{2nuc}$, which has a longer dwell time for ATP, is also weak (Figs. 3a and Supplementary Figure 7a). Taken together, we conclude that the ATP-like density at the catalytic site of $AB_{closed}$ of $V_{3nuc}$ is ADP and phosphate following hydrolysis of ATP.

### Supplementary Figures

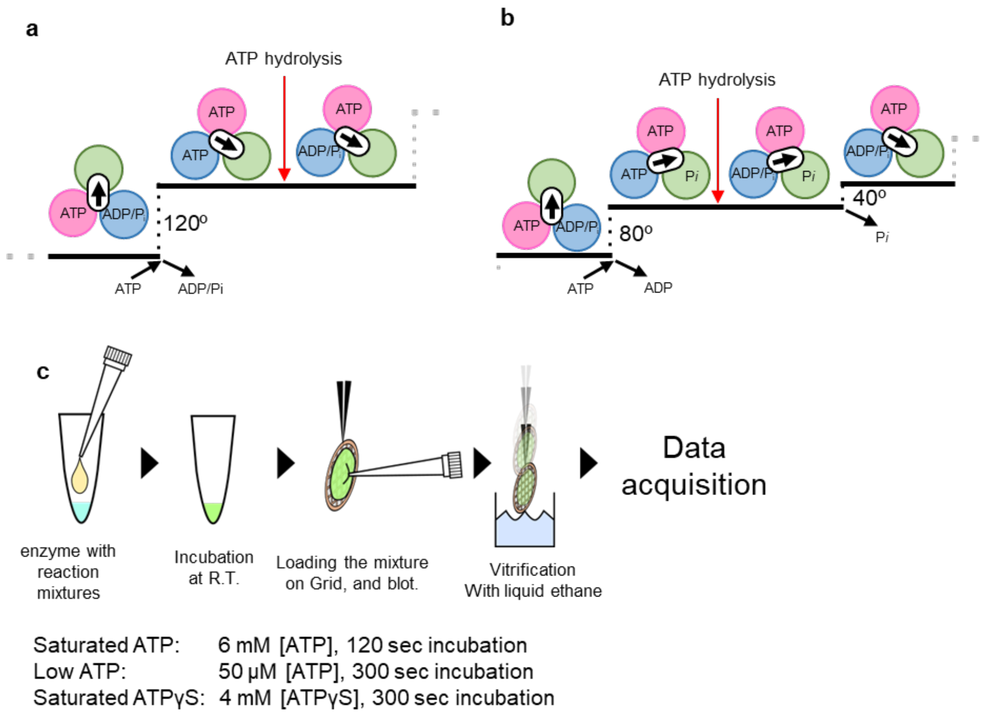


**Supplementary Figure 1. Chemo-mechanical cycle of rotary ATPases.** (**a, b**) The proposed chemo-mechanical cycle of $V_1$-ATPase[3] and $F_1$-ATPase[27]. **(c)** Experimental set up for grid preparation under different reaction conditions.

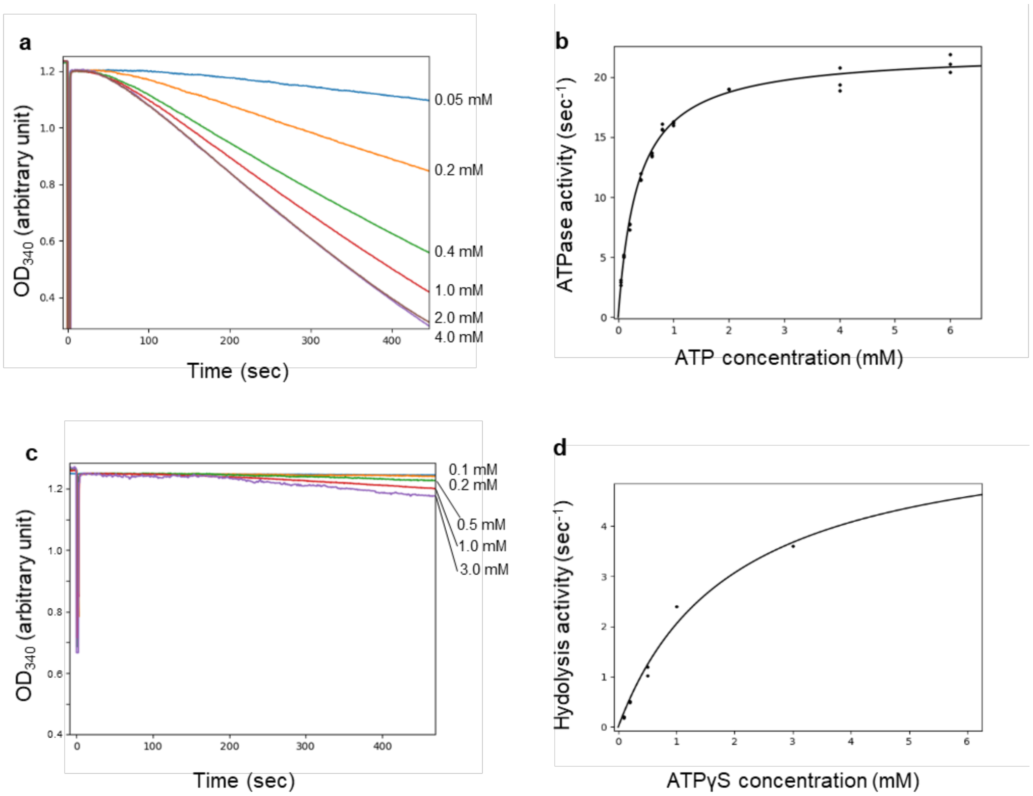


**Supplementary Figure 2. Enzymatic properties of V/A-ATPase for cryo-gird preparation. a**, ATP hydrolysis of nucleotide-free V/A-ATPase incorporated into nanodisc (*nd*-V/A-ATPase) at various ATP concentrations. **b**, *S vs V* plot for ATP hydrolysis by the nucleotide-free *nd*-V/A-ATPase. **c**, ATPγS hydrolysis of nucleotide-free *nd*-V/A-ATPase at various ATPγS concentrations. **d**, *S vs V* plot for ATPγS hydrolysis by the nucleotide-free *nd*-V/A-ATPase. The activities of the *nd*-V/A-ATPase were measured by the enzyme coupling assay described in the Method Details.

#### a: Nucleotide-free

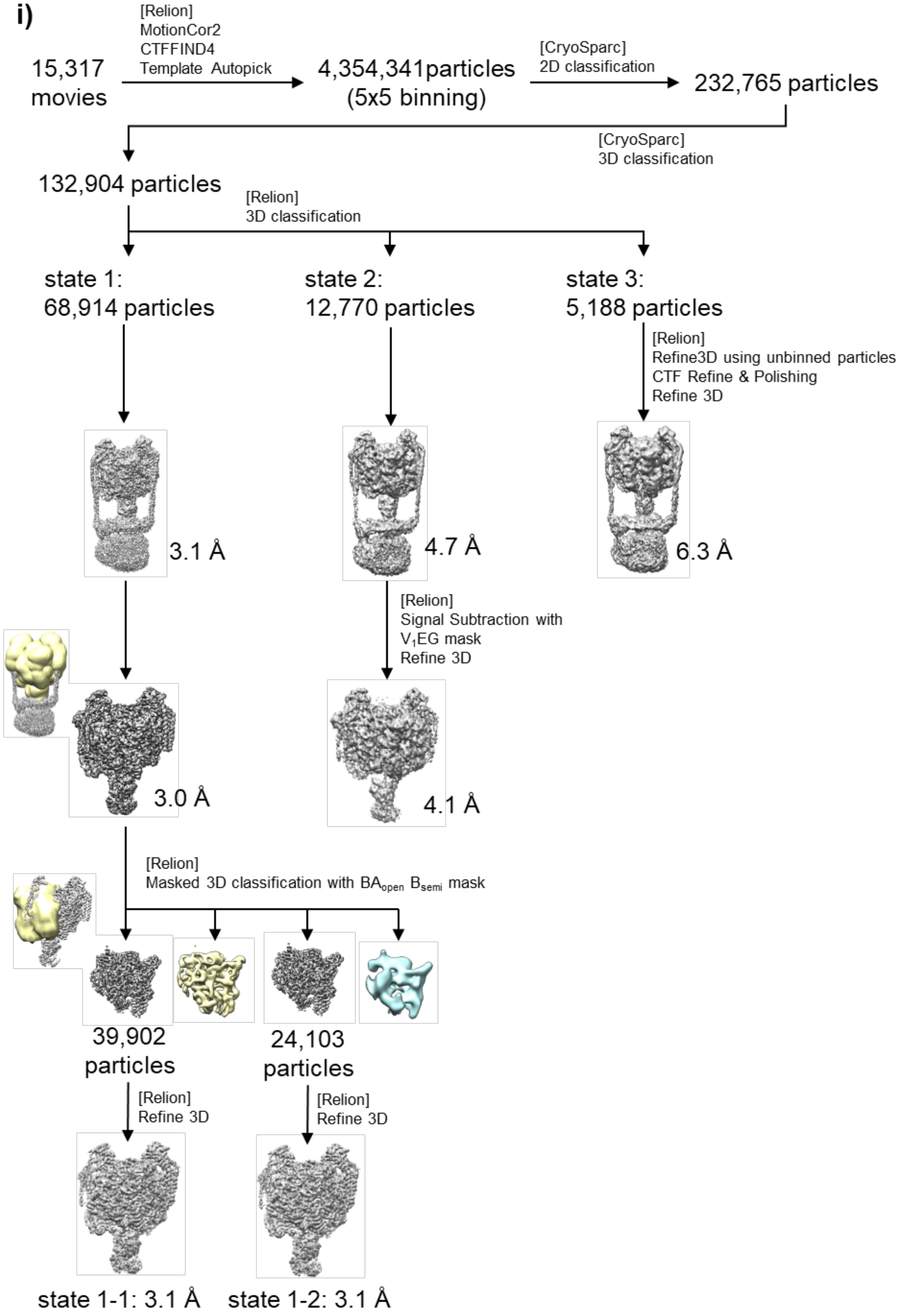

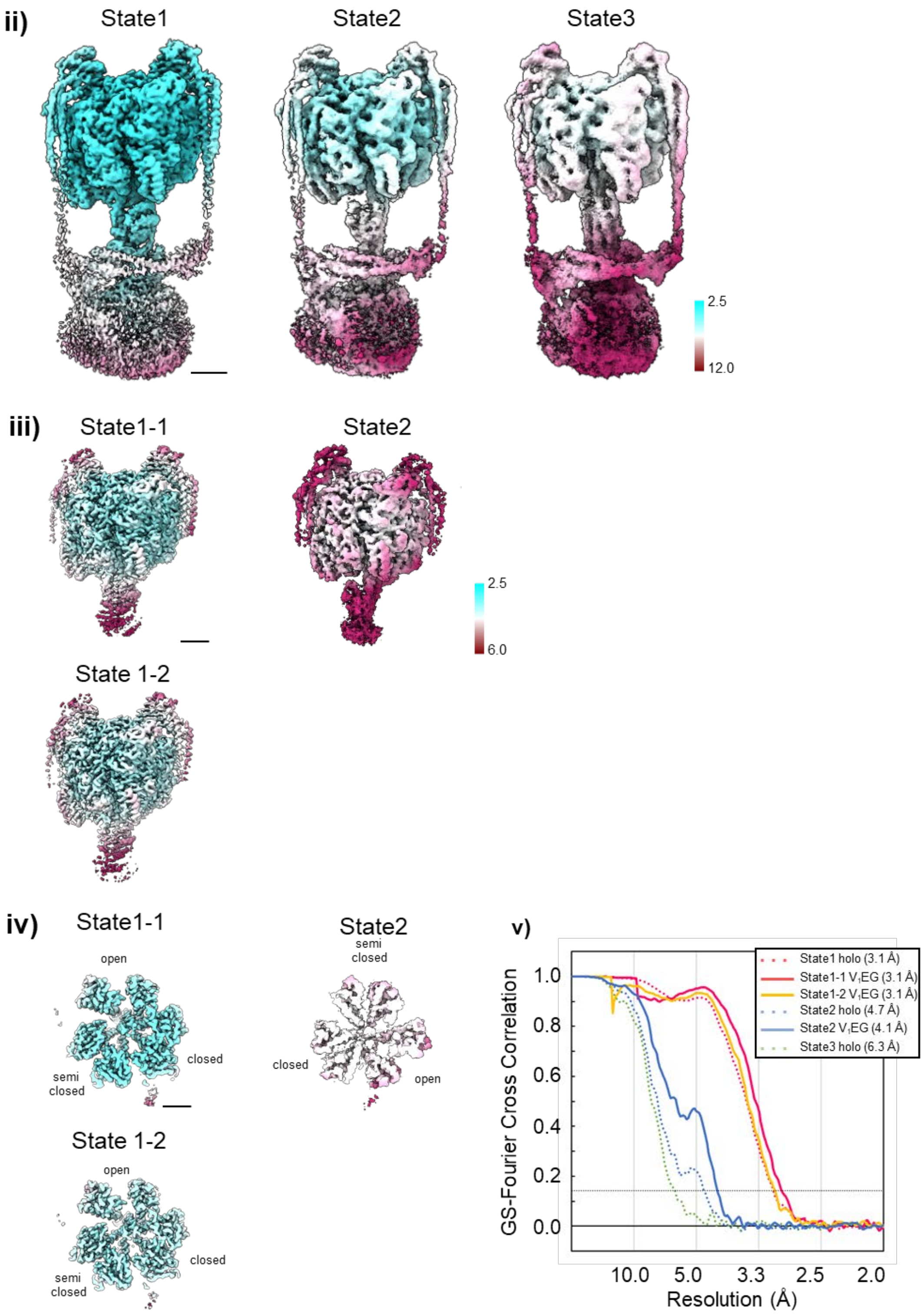
ii)
State1
State2
State3
2.5
12.0
iii)
State1-1
State2
2.5
6.0
State 1-2
iv)
State1-1
open
closed
semi
closed
State2
semi
closed
closed
open
State 1-2
open
closed
semi
closed
v)
State1 holo (3.1 Å)
State1-1 $V_1$EG (3.1 Å)
State1-2 $V_1$EG (3.1 Å)
State2 holo (4.7 Å)
State2 $V_1$EG (4.1 Å)
State3 holo (6.3 Å)
GS-Fourier Cross Correlation
1.0
0.8
0.6
0.4
0.2
0.0
10.0
5.0
3.3
2.5
2.0
Resolution (Å)

#### b: Saturated ATP

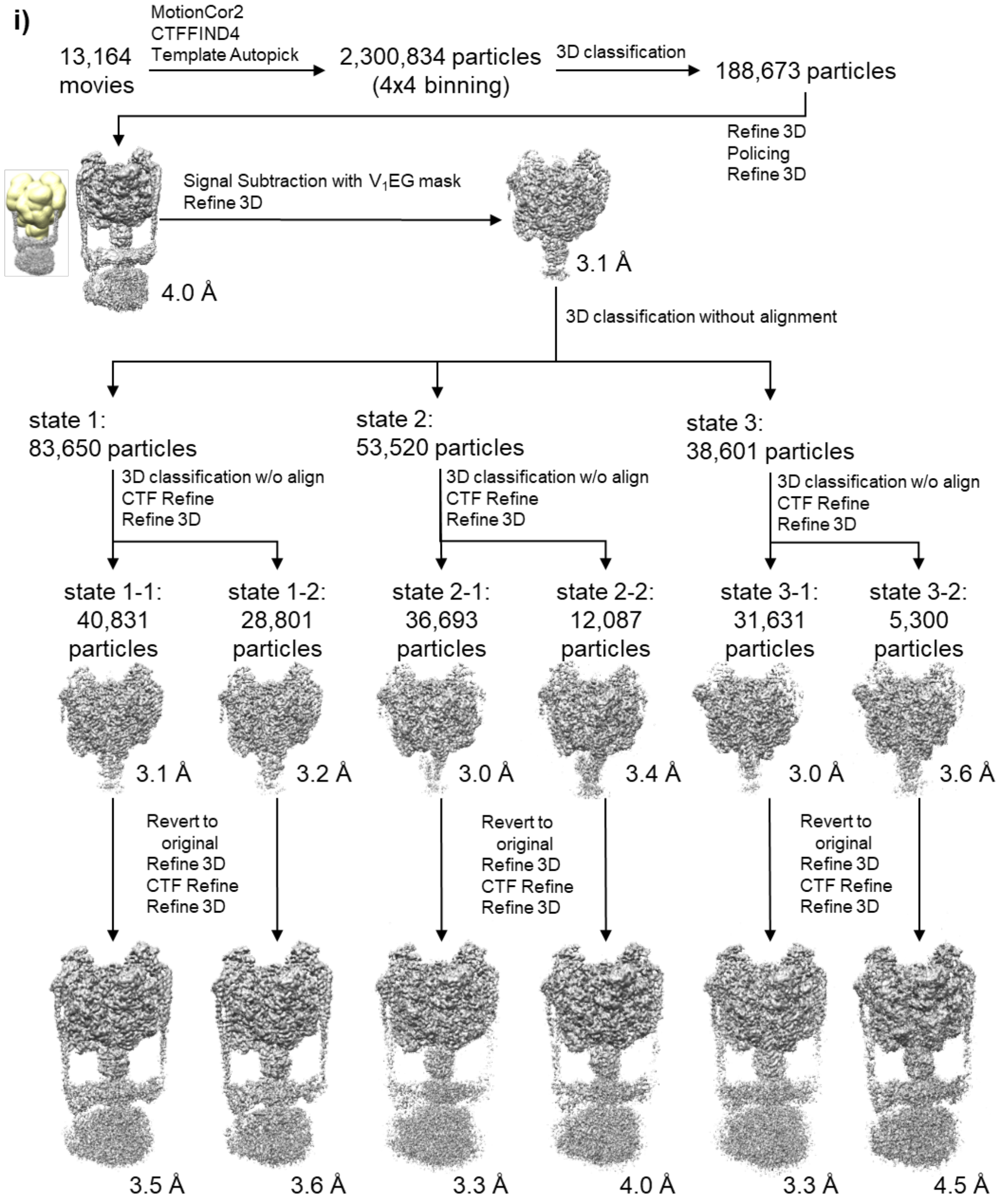

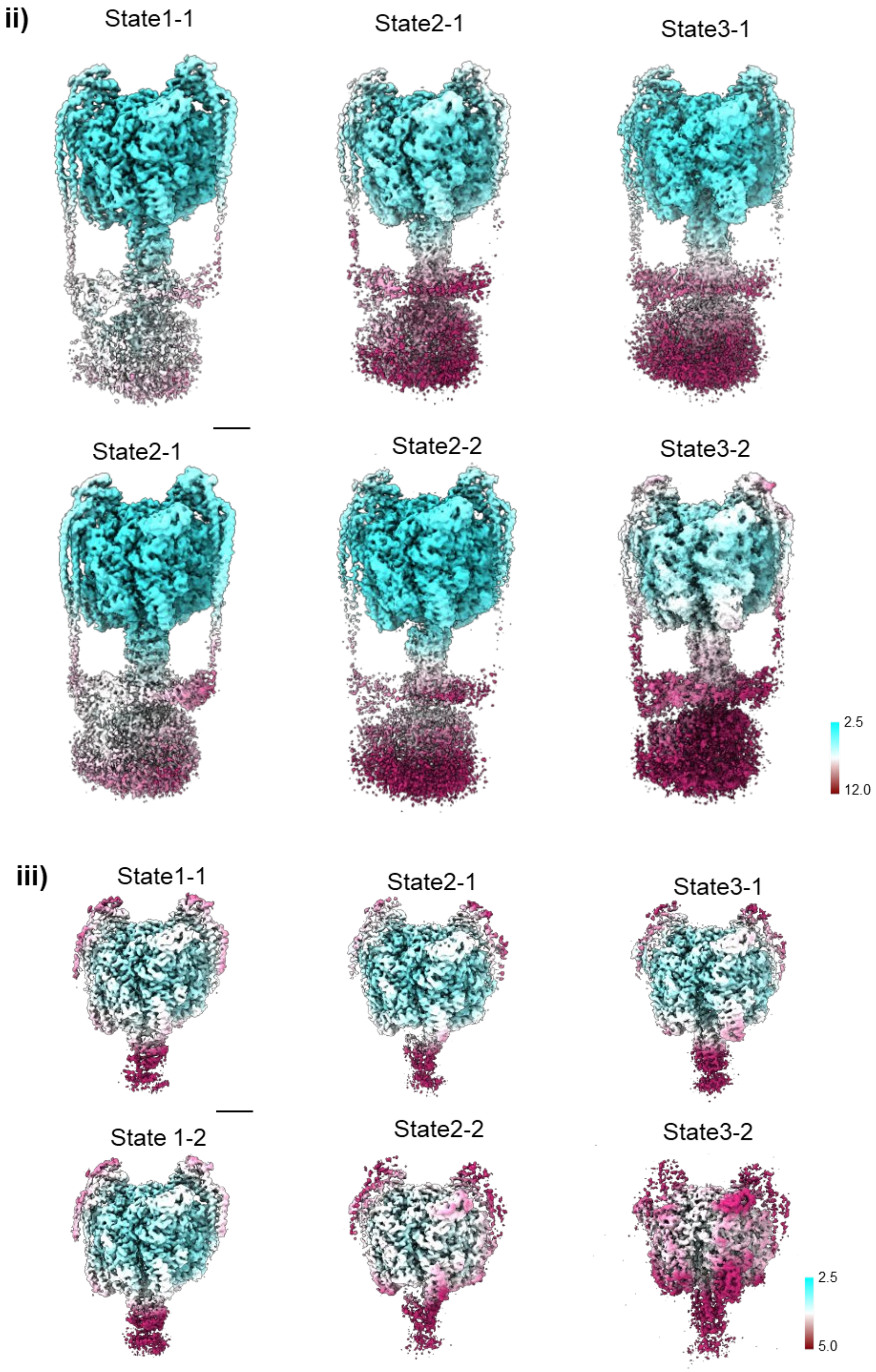

ii)
State1-1
State2-1
State3-1
State2-1
State2-2
State3-2
2.5
12.0
iii)
State1-1
State2-1
State3-1
State 1-2
State2-2
State3-2
2.5
5.0

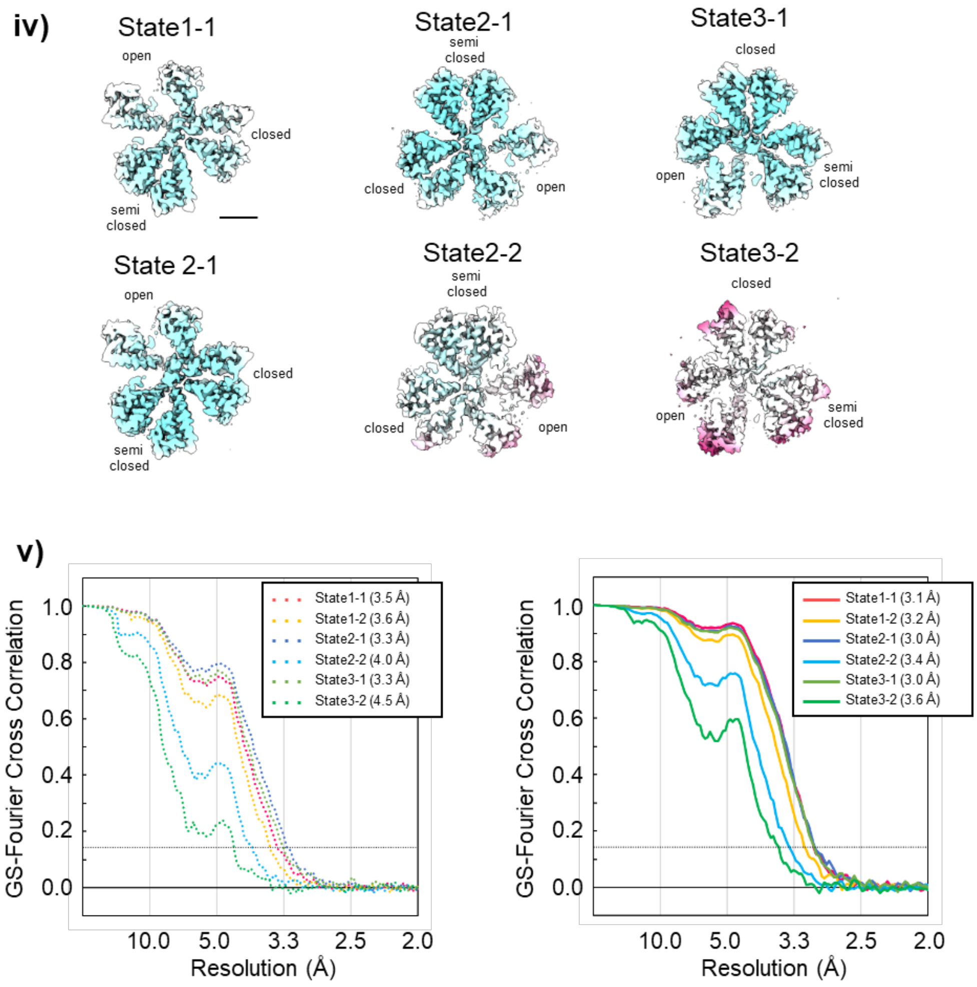
iv)
State1-1
open
closed
semi closed
State2-1
semi closed
closed
open
State3-1
closed
open
semi closed
State 2-1
open
closed
semi closed
State2-2
semi closed
closed
open
State3-2
closed
open
semi closed
v)
GS-Fourier Cross Correlation
1.0
0.8
0.6
0.4
0.2
0.0
10.0
5.0
3.3
2.5
2.0
Resolution (Å)
State1-1 (3.5 Å)
State1-2 (3.6 Å)
State2-1 (3.3 Å)
State2-2 (4.0 Å)
State3-1 (3.3 Å)
State3-2 (4.5 Å)
State1-1 (3.1 Å)
State1-2 (3.2 Å)
State2-1 (3.0 Å)
State2-2 (3.4 Å)
State3-1 (3.0 Å)
State3-2 (3.6 Å)

#### c: Awaiting ATP binding

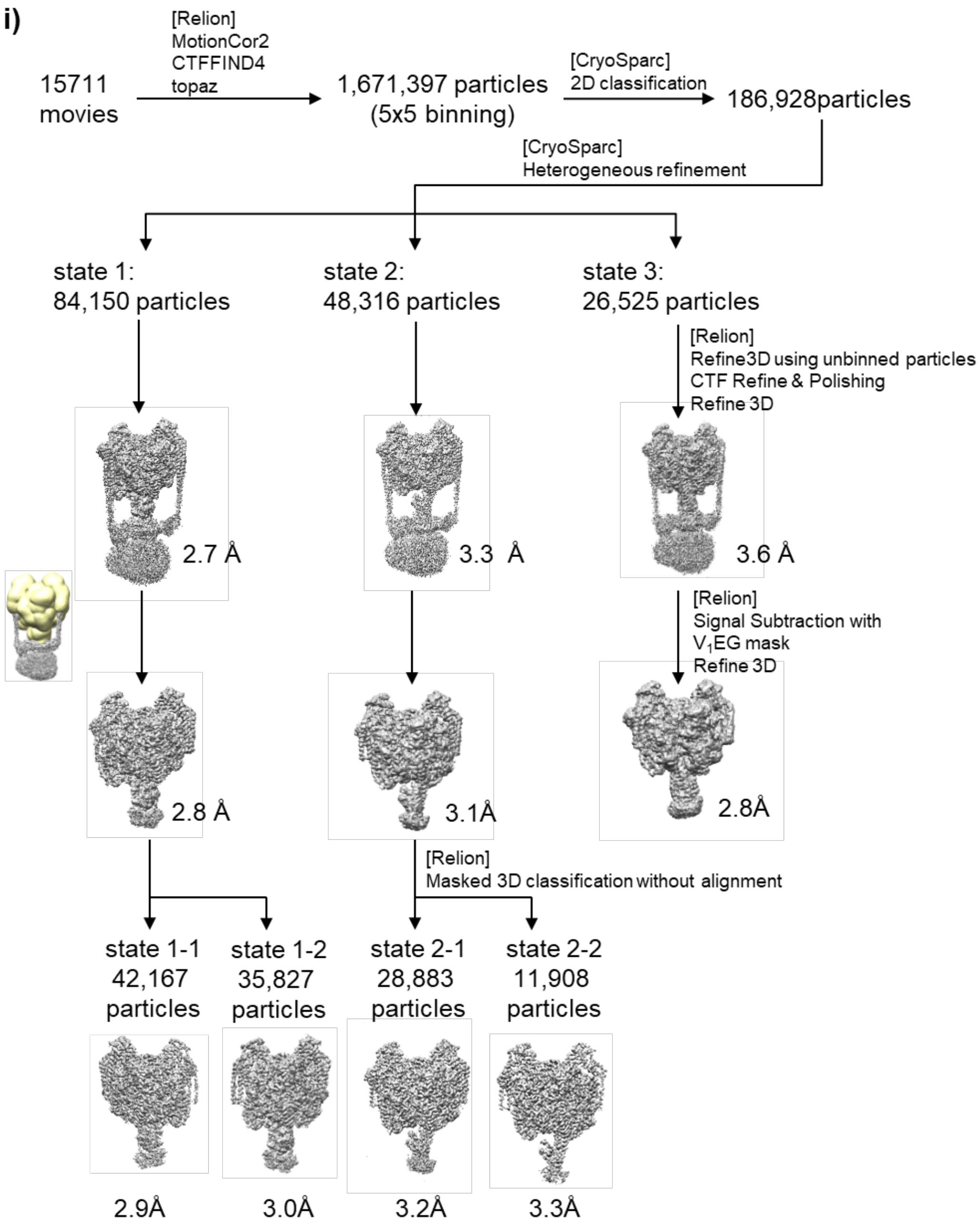

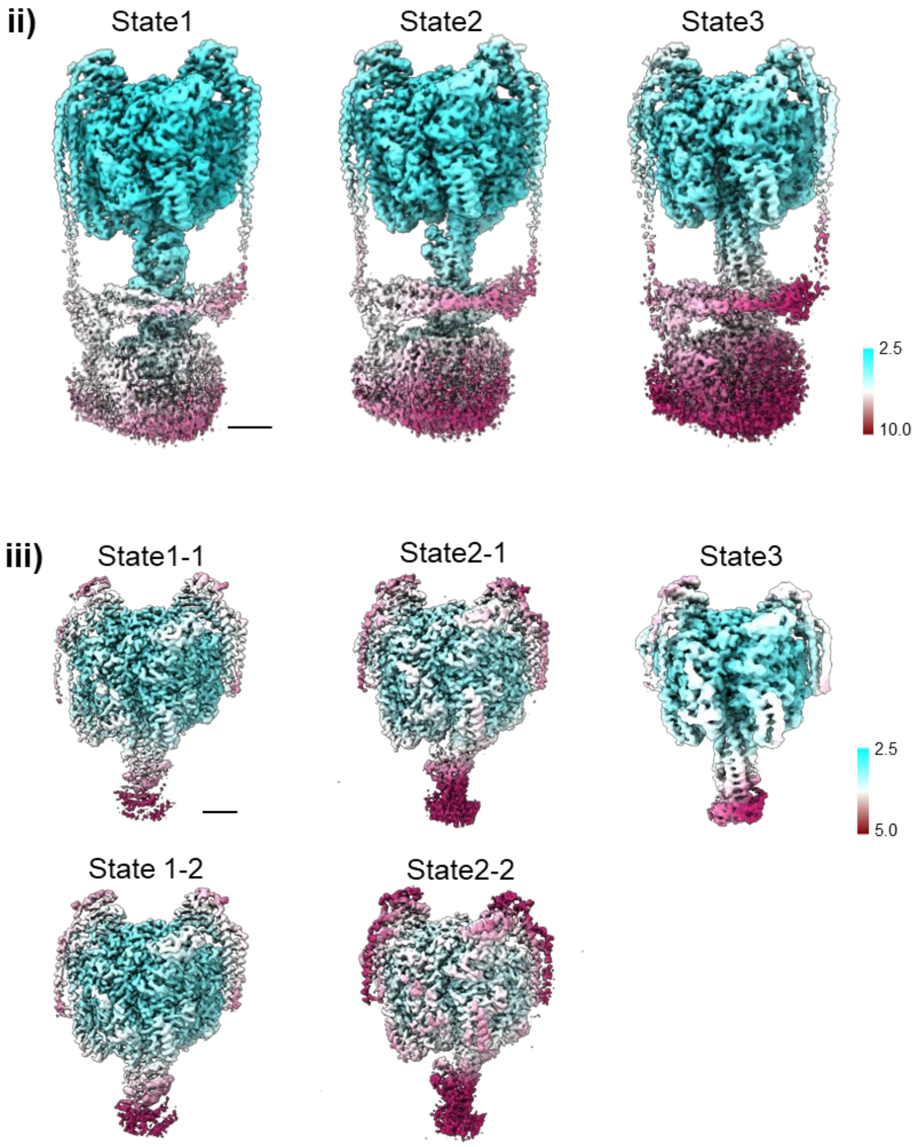
ii)
State1
State2
State3
2.5
10.0
iii)
State1-1
State2-1
State3
2.5
5.0
State 1-2
State2-2

iv)

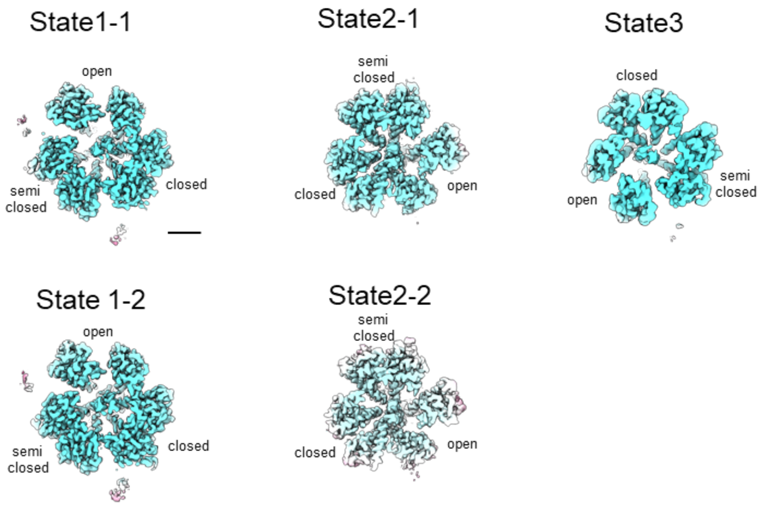
State1-1
open
semi closed
closed
State2-1
semi closed
closed
open
State3
closed
open
semi closed
State 1-2
open
semi closed
closed
State2-2
semi closed
closed
open

v)

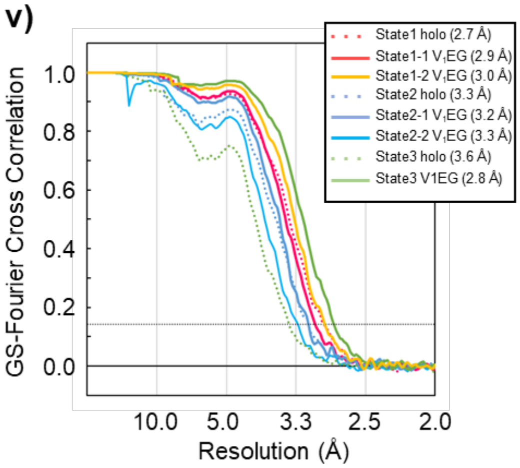
State1 holo (2.7 Å)
State1-1 V1EG (2.9 Å)
State1-2 V1EG (3.0 Å)
State2 holo (3.3 Å)
State2-1 V1EG (3.2 Å)
State2-2 V1EG (3.3 Å)
State3 holo (3.6 Å)
State3 V1EG (2.8 Å)
GS-Fourier Cross Correlation
1.0
0.8
0.6
0.4
0.2
0.0
10.0
5.0
3.3
2.5
2.0
Resolution (Å)

#### d: saturated ATPγS

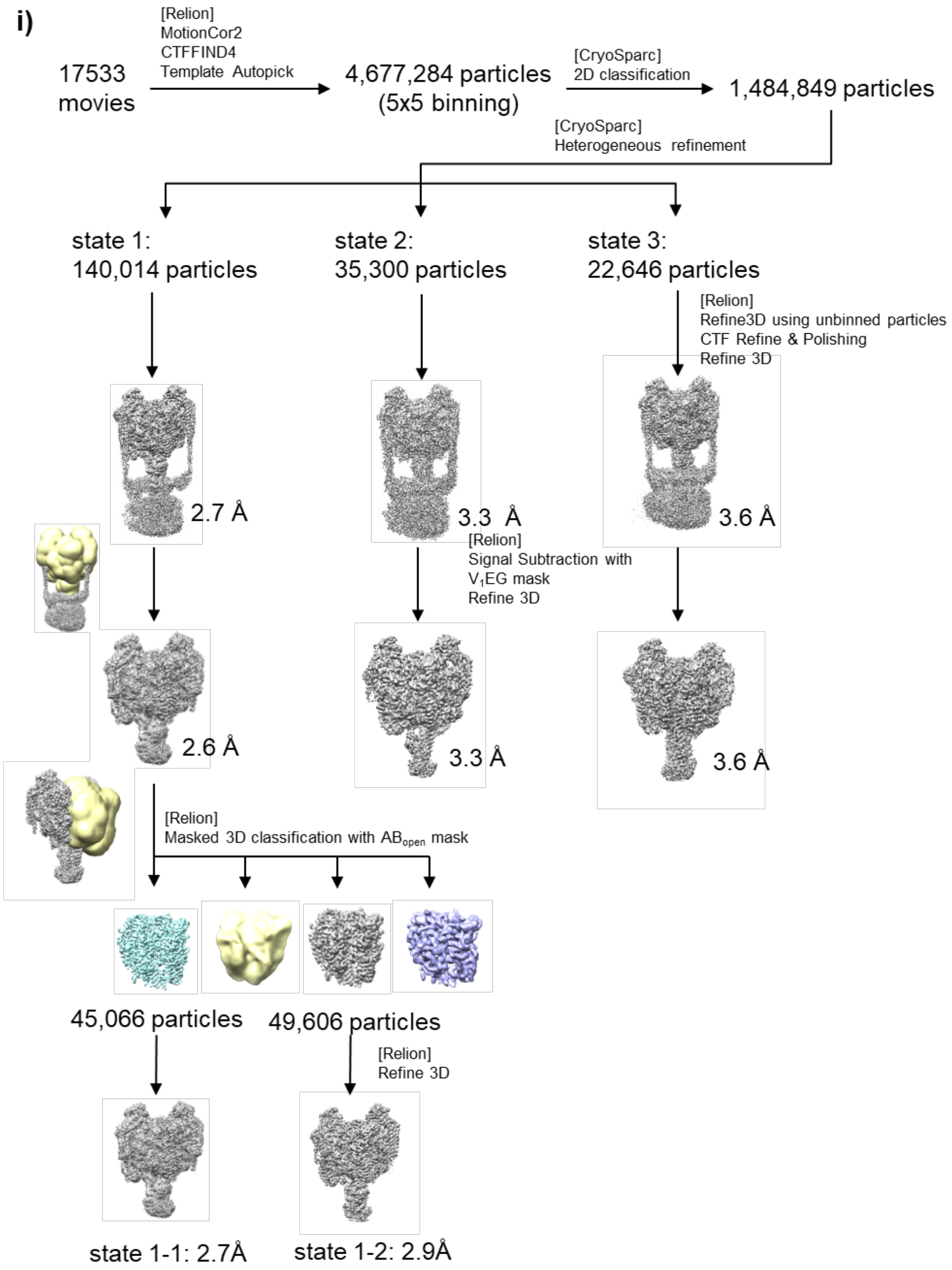

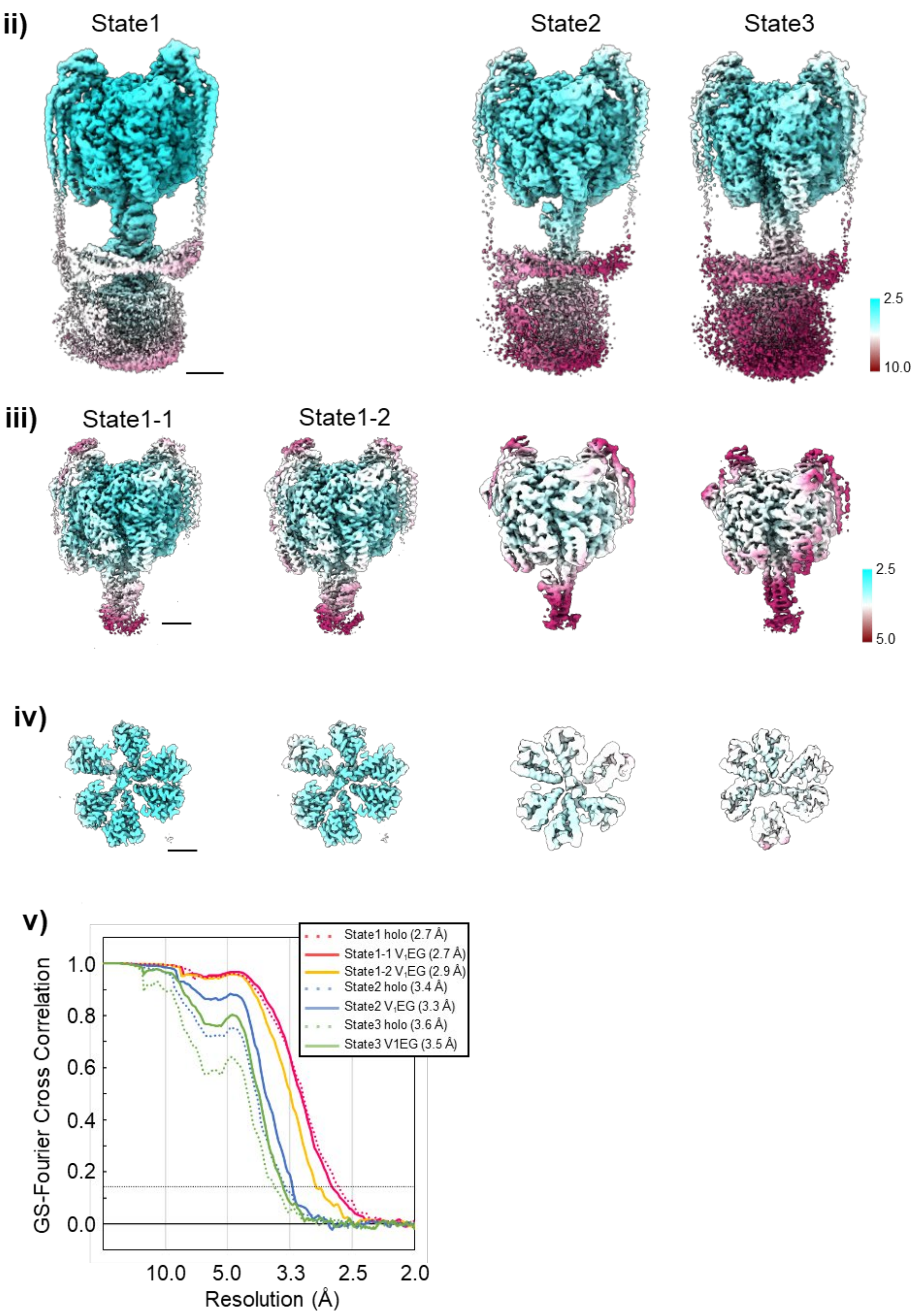

ii)
State1
State2
State3
2.5
10.0
iii)
State1-1
State1-2
2.5
5.0
iv)
v)
State1 holo (2.7 Å)
State1-1 V1EG (2.7 Å)
State1-2 V1EG (2.9 Å)
State2 holo (3.4 Å)
State2 V1EG (3.3 Å)
State3 holo (3.6 Å)
State3 V1EG (3.5 Å)
GS-Fourier Cross Correlation
1.0
0.8
0.6
0.4
0.2
0.0
10.0
5.0
3.3
2.5
2.0
Resolution (Å)

**Supplementary Figure 3. Flow charts describing image acquisition and structural analysis allowing reconstitution of the 3D structures of Nucleotide-free V/A-ATPase (a), at a saturating ATP concentration (b), at ATP waiting condition (c), and at a saturated ATPγS concentration (d).** i) Flow charts describing image acquisition and structural analysis allowing reconstitution of the 3D structures of V/A-ATPase under each condition. ii) Cryo-EM density maps of *holo* V/A-ATPase. iii) Side views of the cryo-EM density maps of $V_1EG$. iv) Cross sections of the $V_1EG$ maps viewed from the cytosolic side. The maps are colored according to local resolution as indicated in the key. v) Gold standard Fourier shell correlation (FSC) curves for *holo* V/A-ATPase (dotted lines) and $V_1EG$ (solid lines), using FSC = 0.143 for resolution criterion. Scale bar is 30 Å.

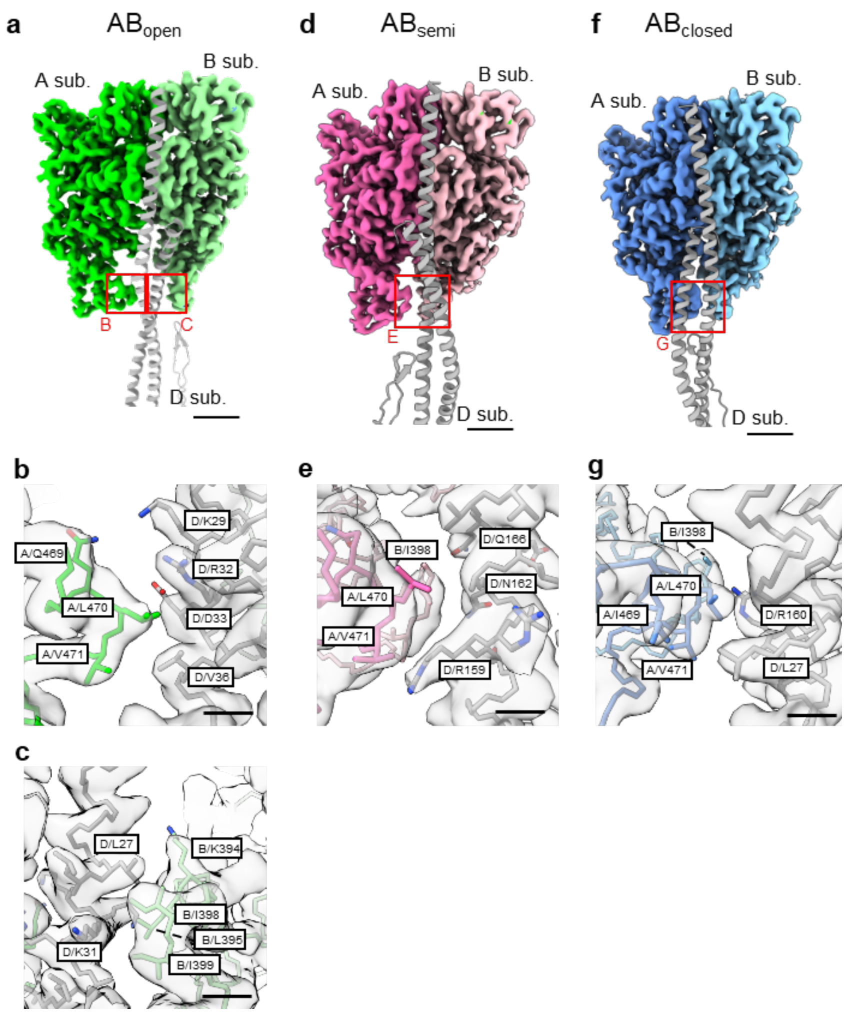


**Suuplementary Figure 4. Interactions of DF with each AB dimer in $V_{nucfree}$. (a, d, f)** The positioning of DF subunits relative to each AB dimer. AB dimers and DF subunits are shown in surface and ribbon representation, respectively. The scale bar is 15 Å. Red squares indicate the interacting interfaces of the CHB and D subunit and are shown enlarged in the lower panels **(b, c, e, g)**. The scale bar is 4 Å.

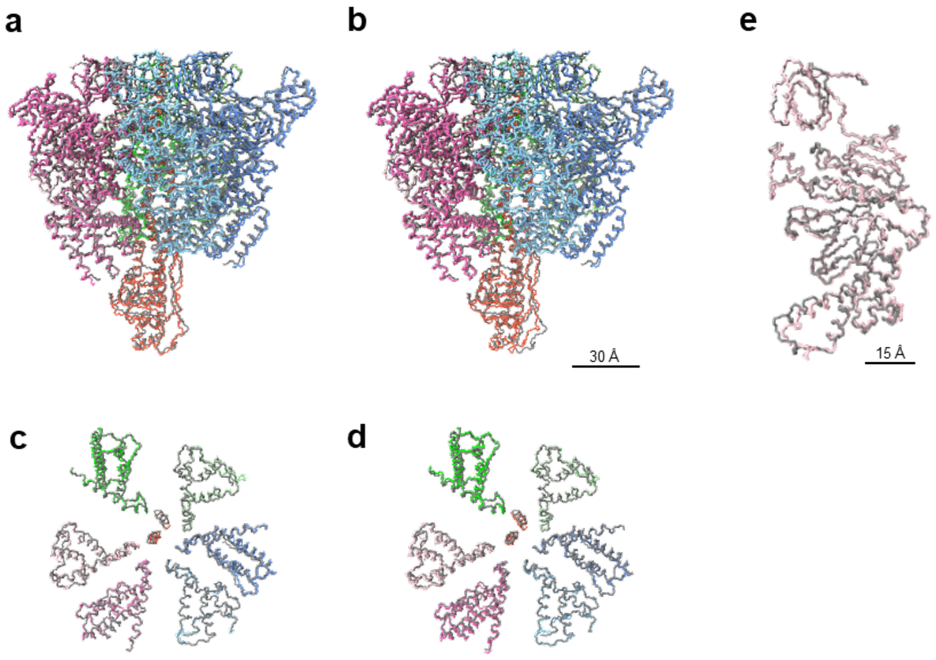


**Supplementary Figure 5. Structural comparison of the $V_{nucfree}$, $V_{3nuc}$, and the ADP inhibited state of V/A-ATPase.** **(a-d)** Comparison of overall and CHB structures of the $V_{nucfree}$ and the inhibited state (PDBID: 6QUM) **(a, c)**, and the $V_{3nuc}$ state1-1 and the inhibited state **(b, d)**. **e,** Comparison of the $B_{semi}$ subunits of $V_{nucfree}$ and $V_{3nuc}$. The structures are superimposed on the β-barrel domain (A subunit 1-70 a.a).

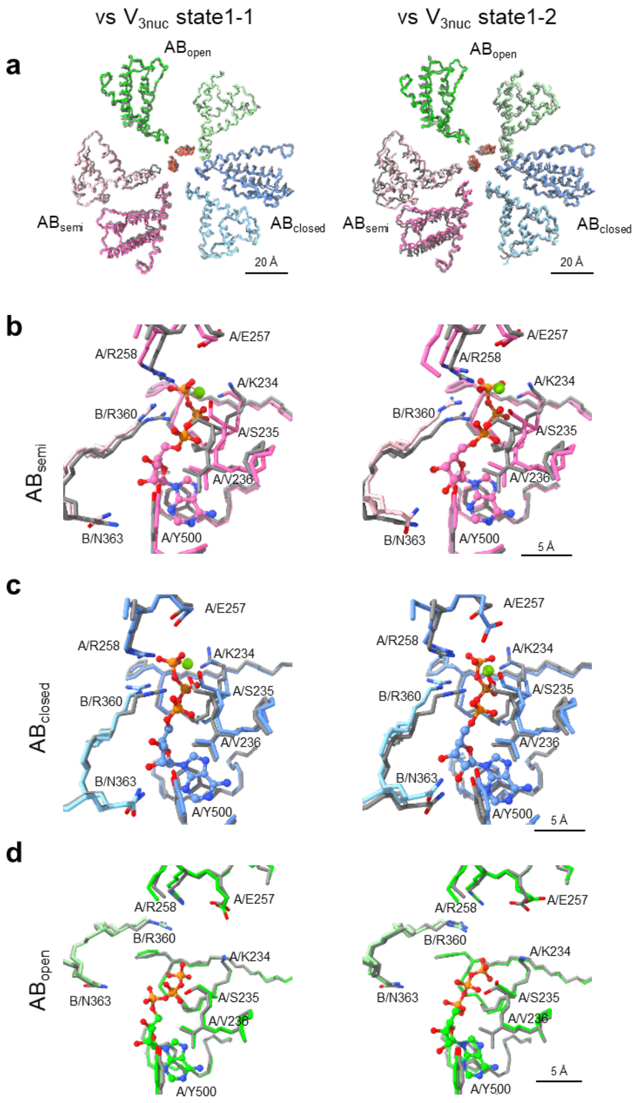


**Supplementary Figure 6. Structural Comparison of $V_{nucfree}$ and $V_{3nuc}$.** Comparison of the three catalytic sites of $V_{3nuc}$ and $V_{nucfree}$. The structures are superimposed on the β barrel domain of the three A subunits (A:1-70 a.a.). **(a)** Comparison of the CHB. **(b-d)** Comparison of the catalytic site of $AB_{semi}$ **(b)**, $AB_{closed}$ **(c)**, and $AB_{open}$ **(d)**. *Left panels;* $V_{nucfree}$ state1 (gray) vs $V_{3nuc}$ state1-1 (colored). *Right panels;* $V_{nucfree}$ state1 (gray) *vs* $V_{3nuc}$ state1-2 (colored).

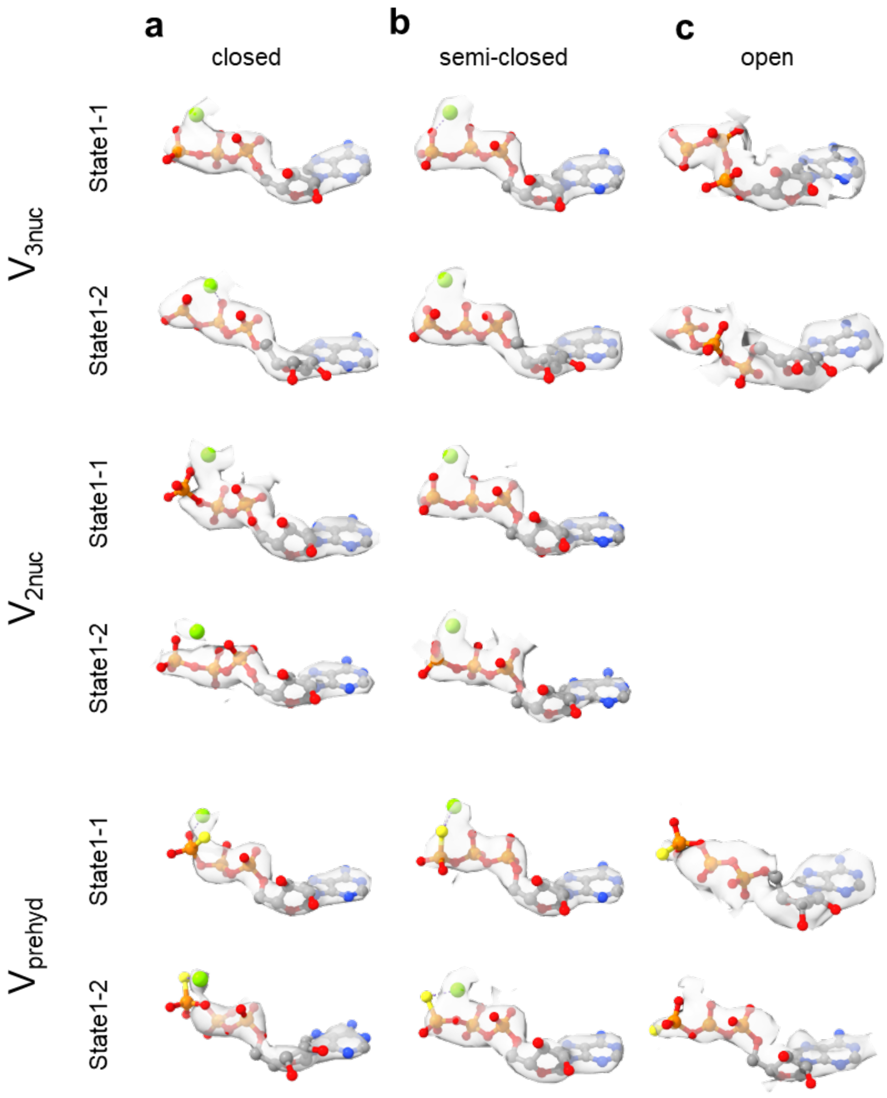


**Supplementary Figure 7. EM density of bound nucleotides at each catalytic site.** The densities corresponding to nucleotides and Mg ions are shown as semi-transparent surfaces. ATP and Mg ions are represented as ball-and-stick and spheres respectively.

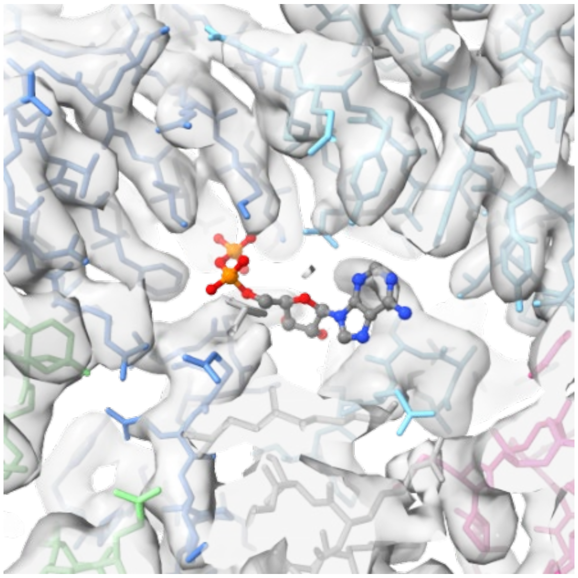

**Supplementary Figure 8. No obvious density for an inhibitory nucleotide.** The previously described structure containing ADP at the tip of the D subunit (PDBID: 6QUM) is fitted into the density map of $V_{3nuc}$ state1-1 from this study. There is no density corresponding to the inhibitory ADP.

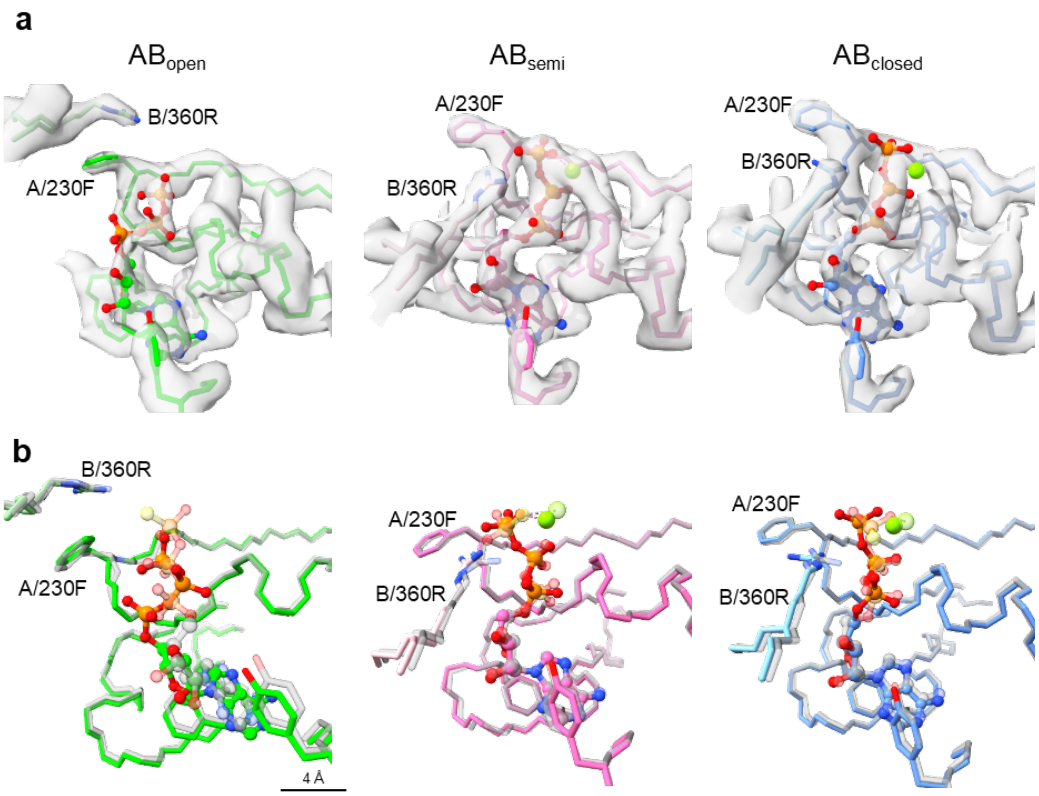


**Supplementary Figure 9. Structural change of A/F230 in $V_{prehyd}$ and $V_{3ATP}$. a**, Magnified views of the three catalytic sites, $AB_{open}$ (*left*), $AB_{semi}$ (*middle*), and $AB_{closed}$ (*right*) of $V_{3nuc}$ state1-1. The chains are shown as sticks. ATP and Mg ions are shown in ball-and-stick and sphere representation, respectively. The density map is represented as a semi-transparent surface. **b**, Comparison of the catalytic sites between $V_{3nuc}$ state1-1 (color) and $V_{prehyd}$ state1-1 (grey).

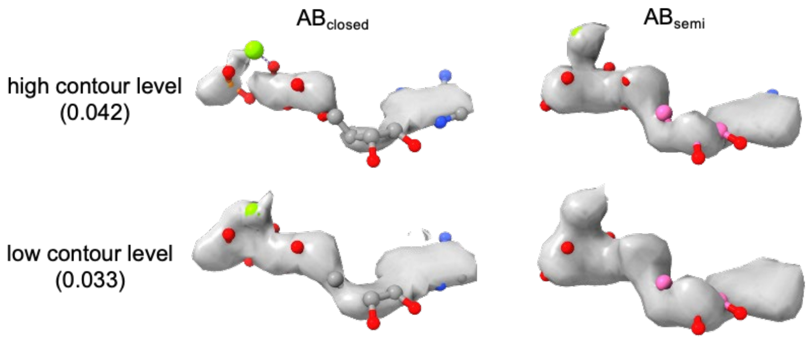


**Supplementary Figure 10. EM density of ATP or ADP and P*i* in AB$_{closed}$ and AB$_{semi}$ of V$_{3nuc}$.** The density corresponding to the bound nucleotides is represented as solid surfaces; AB$_{closed}$ (*left*) and AB$_{semi}$ (*right*). The contour levels are set to relatively high (*upper*) and low (*lower*). The nucleotides and magnesium ions are shown as balls and sticks.

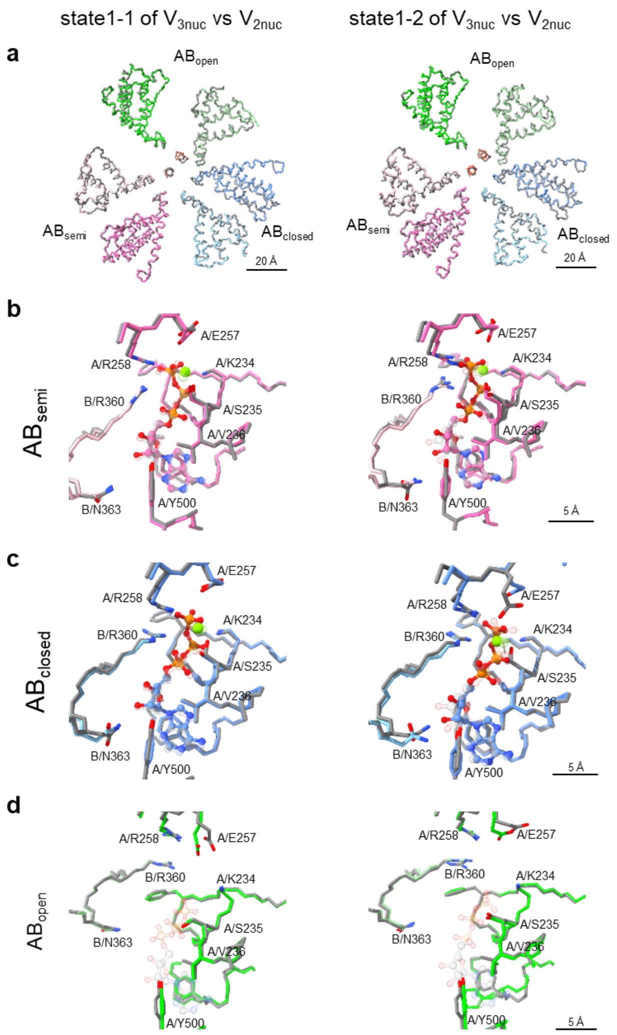


**Supplementary Figure 11. Structural Comparison of $V_{3nuc}$ and $V_{2nuc}$.** Comparison of three catalytic sites between $V_{3nuc}$ and $V_{2nuc}$. The structures are superimposed on the β barrel domain of the three A subunits (A:1-70 a.a.). **(a)** Comparison of the CHB. **(b-d)** Comparison of the catalytic site of $AB_{semi}$ **(b)**, $AB_{closed}$ **(c)**, and $AB_{open}$ **(d)**. *Left panels;* $V_{3nuc}$ state1-1 (gray) vs $V_{2nuc}$ state1-1 (colored). *Right panels;* $V_{3nuc}$ state1-2 (gray) vs $V_{2nuc}$ state1-2 (colored).

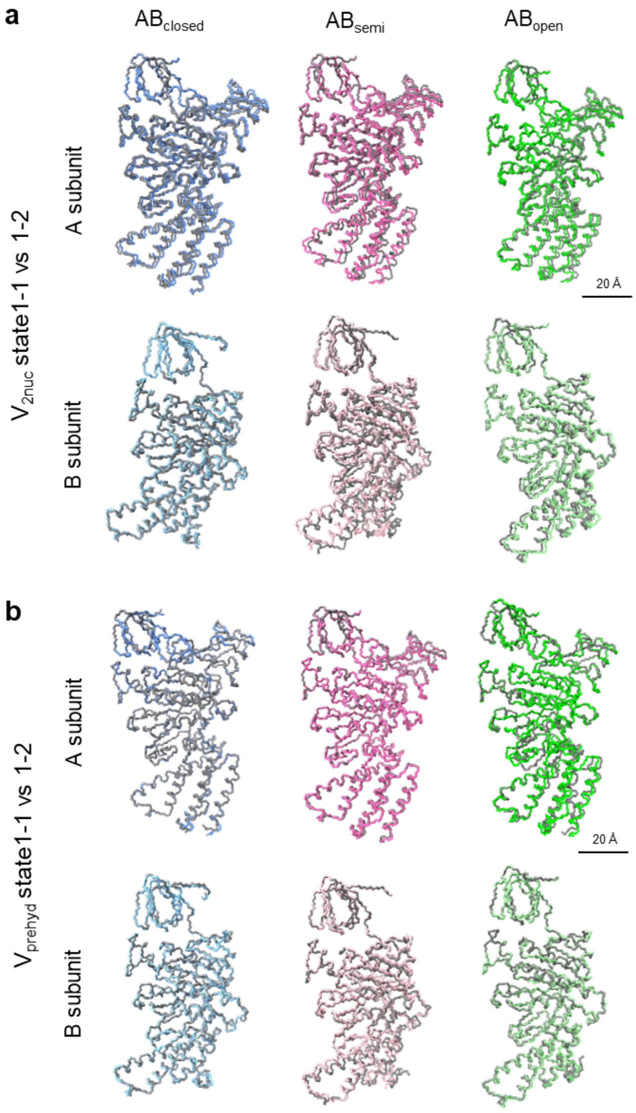


**Supplementary Figure 12. Structural comparisons of each A and B subunit of state1-1 and state 1-2 of $V_{2nuc}$ and $V_{prehyd}$.** Each subunit was superimposed on the β-barrel domain (A subunit 1-70 a.a., B subunit 1-70 a.a.). **a**, Comparisons between state1-1 (gray) and state1-2 (colored) of $V_{2nuc}$. **b**, Comparisons between state1-1 (gray) and state1-2 (colored) of $V_{prehyd}$. The structures are shown as wire models.

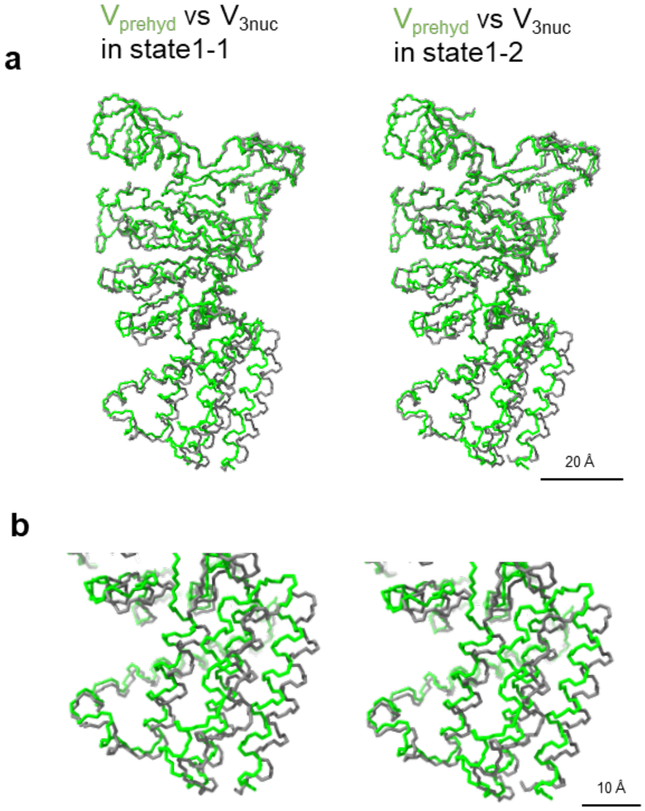


**Supplementary Figure 13. Structural Comparison of $A_{open}$ of $V_{3nuc}$ and $V_{prehyd}$.** The $A_{open}$ subunit in $V_{prehyd}$ state1-2 was compared with that of $V_{3nuc}$ state1-1 and 1-2. The structures are represented as wire models, and are superimposed on the β barrel domain (A:1-70 a.a.). The subunits from $V_{3nuc}$ and $V_{prehyd}$ are colored in gray and green, respectively. **a**, Overall comparison with $V_{3nuc}$ state1-1 (*left*) and state1-2 (*right*). **b**, Enlarged images of CHB shown in (A). RMSD values for the superpositions are summarized in Table S2.

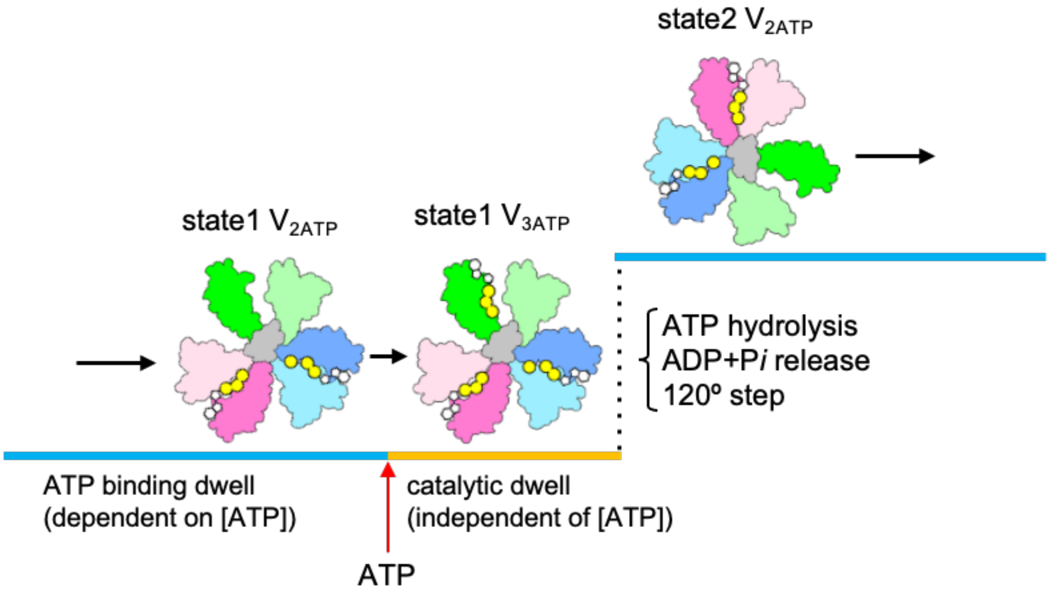


**Supplementary Figure 14. An apparent ATP binding dwell due to awaiting both ATP binding and catalytic events.** Schematic representation of the catalytic cycle of V/A-ATPase during the structural transition from state1 to state2. Following ATP binding, three catalytic processes, ATP hydrolysis in $AB_{semi}$, zippering in $AB_{open}$ and product release in $AB_{closed}$, occur with the 120° step of the DF shaft.

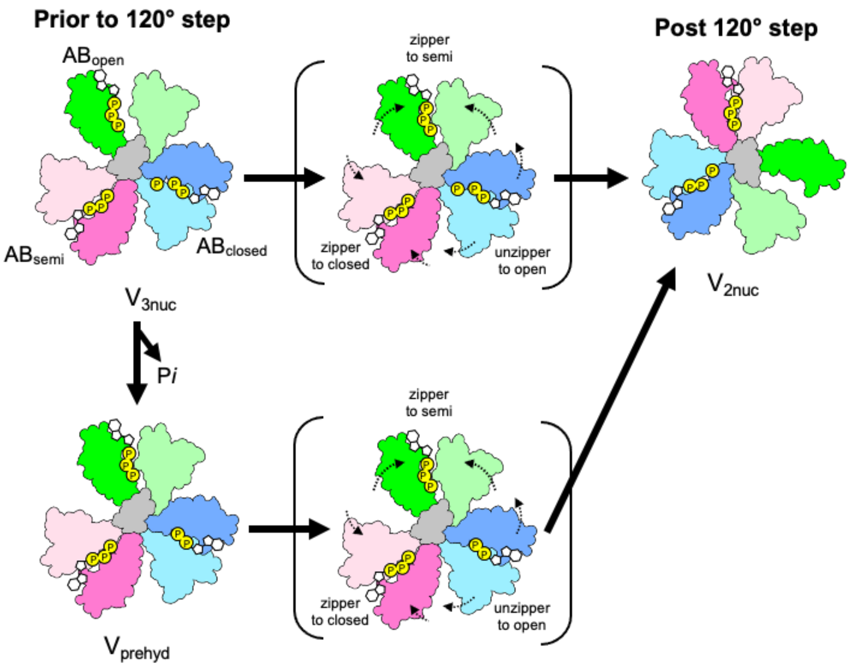


**Supplementary Figure 15. Alternative pathway of V/A-ATPase rotation.** Schematic models of $AB_{open}$, $AB_{semi}$, and $AB_{closed}$ are shown in green, pink, and blue, respectively. The coiled coil region of the D subunit in contact with $A_3B_3$ is shown in grey. The $V_{prehyd}$ corresponds to the structure in which phosphate is released from $AB_{closed}$. The catalytic events of the three AB dimers of $V_{prehyd}$ occur simultaneously as in $V_{3nuc}$, resulting in structural transition of $V_{prehyd}$ to $V_{2nuc}$.

**Supplementary Table 1. Number of bound nucleotides of V/A-ATPase with or without nucleotide removal treatment.**

| | WT V/A-ATPase | | TSSA V/A-ATPase | |
|---|---|---|---|---|
| | non-treated | treated | non-treated | Treated |
| ATP[1] | 1.9 | 1.4 | 0.27 | <0.01 |
| ADP[1] | 0.73 | 0.3 | <0.01 | <0.01 |
| ATPase activity ($s^{-1}$) | <1 | 5 | 45 | 51 |

[1]Number of binding nucleotides per enzyme.

**Supplementary Table 2. RMSD values for A subunits of each state.** The subunits were superimposed on the β-barrel domain (1-70 a.a), then the values for the back bone (444-573 a.a) were calculated using UCSF Chimera software. The lowest value in each row is indicated in bold.

(Å)

| | 444-573 (bb only) | $V_{3nuc}$ state1-1 | $V_{3nuc}$ state1-2 | $V_{2nuc}$ state1-1 | $V_{2nuc}$ state1-2 | $V_{prehyd}$ state1-1 | $V_{prehyd}$ state1-2 | $V_{nucfree}$ State1-1 | $V_{nucfree}$ State1-2 |
|---|---|---|---|---|---|---|---|---|---|
| $A_{open}$ | $V_{3nuc}$ state1-1 | | 1.603 | **0.534** | 1.721 | 1.633 | 4.094 | 0.575 | 1.579 |
| | $V_{3nuc}$ state1-2 | 1.603 | | 1.283 | **0.485** | 1.550 | 3.962 | 1.440 | 0.544 |
| | $V_{2nuc}$ state1-1 | **0.534** | 1.283 | | 1.329 | 1.455 | 4.042 | 0.423 | 1.253 |
| | $V_{2nuc}$ state1-2 | 1.721 | **0.485** | 1.329 | | 1.549 | 4.104 | 1.485 | 0.389 |
| | $V_{prehyd}$ state1-1 | 1.633 | 1.550 | **1.455** | 1.549 | | 3.480 | 1.614 | 1.522 |
| | $V_{prehyd}$ state1-2 | 4.094 | 3.962 | 4.042 | 4.104 | **3.48** | | 4.305 | 4.138 |
| | $V_{nucfree}$ state1-1 | 0.575 | 1.440 | **0.423** | 1.485 | 1.614 | 4.305 | | 1.278 |
| | $V_{nucfree}$ state1-2 | 1.579 | 0.544 | 1.253 | **0.389** | 1.522 | 4.138 | 1.278 | |
| $A_{semi}$ | $V_{3nuc}$ state1-1 | | 2.306 | **0.369** | 2.603 | 1.576 | 1.990 | 0.602 | 2.041 |
| | $V_{3nuc}$ state1-2 | 2.306 | | 2.081 | **0.383** | 1.855 | 1.365 | 1.936 | 0.548 |
| | $V_{2nuc}$ state1-1 | **0.369** | 2.081 | | 2.390 | 1.413 | 1.783 | 0.464 | 1.789 |
| | $V_{2nuc}$ state1-2 | 2.603 | **0.383** | 2.39 | | 2.092 | 1.524 | 2.206 | 0.720 |
| | $V_{prehyd}$ state1-1 | 1.576 | 1.855 | 1.413 | 2.092 | | **0.844** | 1.530 | 1.759 |
| | $V_{prehyd}$ state1-2 | 1.99 | 1.365 | 1.783 | 1.524 | **0.844** | | 1.733 | 1.363 |
| | $V_{nucfree}$ state1-1 | 0.602 | 1.936 | **0.464** | 2.206 | 1.503 | 1.733 | | 1.664 |
| | $V_{nucfree}$ state1-2 | 2.041 | **0.548** | 1.789 | 0.720 | 1.759 | 1.363 | 1.664 | |
| $A_{closed}$ | $V_{3nuc}$ state1-1 | | 1.714 | **0.278** | 1.691 | 1.339 | 1.403 | 0.578 | 1.685 |
| | $V_{3nuc}$ state1-2 | 1.714 | | 1.642 | **0.292** | 2.262 | 2.095 | 1.496 | 0.457 |
| | $V_{2nuc}$ state1-1 | **0.278** | 1.642 | | 1.632 | 1.307 | 1.359 | 0.487 | 1.594 |
| | $V_{2nuc}$ state1-2 | 1.691 | **0.292** | 1.632 | | 2.289 | 2.119 | 1.472 | 0.382 |
| | $V_{prehyd}$ state1-1 | 1.339 | 2.262 | 1.307 | 2.289 | | **0.381** | 1.428 | 2.325 |
| | $V_{prehyd}$ state1-2 | 1.403 | 2.095 | 1.359 | 2.119 | **0.381** | | 1.395 | 2.155 |
| | $V_{nucfree}$ state1-1 | 0.578 | 1.496 | **0.487** | 1.472 | 1.428 | 1.395 | | 1.413 |
| | $V_{nucfree}$ state1-2 | 1.685 | 0.457 | 1.594 | **0.382** | 2.325 | 2.155 | 1.413 | |

**Supplementary Table 3. RMSD values for B subunits of each state.** The subunits were superimposed on the β-barrel domain (1-70 a.a), then the values for the back bone (374-446 a.a) were calculated using UCSF Chimera software. The lowest value in each row is indicated in bold.

(Å)

| | 374-446 (bb only) | $V_{3nuc}$ state1-1 | $V_{3nuc}$ state1-2 | $V_{2nuc}$ state1-1 | $V_{2nuc}$ state1-2 | $V_{prehyd}$ state1-1 | $V_{prehyd}$ state1-2 | $V_{nucfree}$ state1-1 | $V_{nucfree}$ state1-2 |
|---|---|---|---|---|---|---|---|---|---|
| $B_{open}$ | $V_{3nuc}$ state 1-1 | | 1.531 | **0.314** | 1.310 | 1.168 | 1.200 | 0.346 | 1.309 |
| | $V_{3nuc}$ state 1-2 | 1.531 | | 1.607 | **0.477** | 2.118 | 1.786 | 1.714 | 0.439 |
| | $V_{2nuc}$ state1-1 | **0.314** | 1.607 | | 1.356 | 1.197 | 1.280 | 0.333 | 1.337 |
| | $V_{2nuc}$ state1-2 | 1.31 | **0.477** | 1.356 | | 1.967 | 1.682 | 1.487 | 0.325 |
| | $V_{prehyd}$ state1-1 | 1.168 | 2.118 | 1.197 | 1.967 | | **0.535** | 1.257 | 2.037 |
| | $V_{prehyd}$ state1-2 | 1.2 | 1.786 | 1.28 | 1.682 | **0.535** | | 1.357 | 1.770 |
| | $V_{nucfree}$ state1-1 | 0.346 | 1.714 | **0.333** | 1.487 | 1.257 | 1.357 | | 1.488 |
| | $V_{nucfree}$ state1-2 | 1.309 | 0.439 | 1.337 | **0.325** | 2.037 | 1.770 | 1.488 | |
| $B_{semi}$ | $V_{3nuc}$ state1-1 | | 3.647 | **0.517** | 3.594 | 1.992 | 3.268 | 0.803 | 3.240 |
| | $V_{3nuc}$ state1-2 | 3.647 | | 3.561 | **0.526** | 2.755 | 2.161 | 3.906 | 1.014 |
| | $V_{2nuc}$ state1-1 | **0.517** | 3.561 | | 3.512 | 1.885 | 3.142 | 0.606 | 3.109 |
| | $V_{2nuc}$ state1-2 | 3.594 | **0.526** | 3.512 | | 2.732 | 2.295 | 3.840 | 0.757 |
| | $V_{prehyd}$ state1-1 | 1.992 | 2.755 | 1.885 | 2.732 | | 2.096 | 2.240 | 2.465 |
| | $V_{prehyd}$ state1-2 | 3.268 | 2.161 | 3.142 | 2.295 | **2.096** | | 3.464 | 2.206 |
| | $V_{nucfree}$ state1-1 | 0.803 | 3.906 | **0.606** | 3.840 | 2.240 | 3.464 | | 3.405 |
| | $V_{nucfree}$ state1-2 | 3.240 | 1.014 | 3.109 | **0.757** | 2.465 | 2.206 | 3.405 | |
| $B_{closed}$ | $V_{3nuc}$ state1-1 | | 1.032 | **0.352** | 1.152 | 1.144 | 1.345 | 0.468 | 1.045 |
| | $V_{3nuc}$ state1-2 | 1.032 | | 1.155 | **0.369** | 1.364 | 1.240 | 1.172 | 0.419 |
| | $V_{2nuc}$ state1-1 | **0.352** | 1.155 | | 1.219 | 1.228 | 1.427 | 0.419 | 1.073 |
| | $V_{2nuc}$ state1-2 | 1.152 | **0.369** | 1.219 | | 1.451 | 1.266 | 1.244 | 0.422 |
| | $V_{prehyd}$ state1-1 | 1.144 | 1.364 | 1.228 | 1.451 | | **0.542** | 1.342 | 1.546 |
| | $V_{prehyd}$ state1-2 | 1.345 | 1.24 | 1.427 | 1.266 | **0.542** | | 1.495 | 1.409 |
| | $V_{nucfree}$ state1-1 | 0.468 | 1.172 | **0.419** | 1.244 | 1.342 | 1.495 | | 1.096 |
| | $V_{nucfree}$ state1-2 | 1.045 | **0.419** | 1.073 | 0.422 | 1.546 | 1.409 | 1.096 | |

**Supplementary Table 4. Cryo-EM data collection, refinement and validation statistics for nucleotide-free V/A-ATPase.**

| | State1 | | | State2 | | State3 |
|---|---|---|---|---|---|---|
| | $V_oV_1$ | $V_1$EG state1-1 | $V_1$EG state1-2 | $V_oV_1$ | $V_1$EG | $V_oV_1$ |
| EMDB ID | 31841 | 31842 | 31843 | 31844 | 31845 | 31846 |
| PDB ID | | 7VAI | 7VAJ | | 7VAK | |
| **Data collection and processing** | | | | | | |
| Magnification | 81,000x | | | | | |
| EM & Voltage (kV) | Titan Krios, 300 | | | | | |
| Total dose (e-/Å$^2$) | 50 | | | | | |
| Pixel size (Å/pix) | 0.88 | | | | | |
| Defocus range (μm) | -0.8 to -2.0 | | | | | |
| Symmetry imposed | C1 | | | | | |
| Initial particle # | 4,354,341 | | | | | |
| Final Particle # | 68,290 | 39,902 | 24,103 | 12,770 | 12,770 | 5,188 |
| Map resolution (Å) | 3.1 | 3.1 | 3.1 | 4.7 | 4.1 | 6.3 |
| FSC threshold | 0.143 | | | | | |
| **Refinement** | | | | | | |
| Initial model used (PDBID) | - | 6QUM | This study | - | This study | - |
| Model resolution (Å) | - | 3.1 (masked) | 3.1 (masked) | - | 4.3 (masked) | - |
| FSC threshold | - | 0.5 | 0.5 | - | 0.5 | - |
| Model composition | | | | | | |
| Nonhydrogen atoms | - | 29,462 | 29,462 | - | 29,462 | - |
| Protein residues | - | 3,788 | 3,788 | | 3,788 | - |
| Ligands | - | 0 | 0 | | 0 | - |
| R.m.s deviations | | | | | | |
| Bond length (Å) | - | 0.011 | 0.10 | | 0.003 | - |
| Bond Angles (°) | - | 0.814 | 0.779 | | 0.549 | - |
| **Validation** | | | | | | |
| MolProbity score | - | 1.71 | 1.74 | | 1.64 | - |
| EMRinger score | | 3.41 | 2.98 | | 1.59 | - |
| Clashscore | - | 6.94 | 7.27 | | 8.89 | - |
| Rotamer outlier (%) | - | 0.03 | 0.05 | | 0.00 | - |
| CaBALM outlierl (%) | | 2.41 | 2.41 | | 2.01 | - |
| Ramachandran plot | | | | | | |
| Favored (%) | - | 95.32 | 95.14 | | 97.05 | - |
| Allowed (%) | - | 4.65 | 4.81 | | 2.95 | - |
| Disallowed (%) | - | 0.03 | 0.05 | | 0.00 | - |

**Supplementary Table 5-1. Cryo-EM data collection, refinement and validation statistics for V/A-ATPase obtained under 6 mM ATP condition.**

| | State1-1 | | State1-2 | |
|---|---|---|---|---|
| | $V_oV_1$ | $V_1$EG | $V_oV_1$ | $V_1$EG |
| EMDB ID | 31847 | 31849 | 31848 | 31850 |
| PDB ID | | 7VAL | | 7VAM |
| **Data collection and processing** | | | | |
| Magnification | 81,000x | | | |
| EM & Voltage (kV) | Titan Krios, 300 | | | |
| Total dose (e-/Å$^2$) | 50 | | | |
| Pixel size (Å/pix) | 0.88 | | | |
| Defocus range (μm) | -0.8 to -2.0 | | | |
| Symmetry imposed | C1 | | | |
| Initial particle # | 2,300,834 | | | |
| Final Particle # | 40,831 | | 28,801 | |
| Map resolution (Å) | 3.5 | 3.1 | 3.6 | 3.2 |
| FSC threshold | 0.143 | | | |
| **Refinement** | | | | |
| Initial model used (PDBID) | - | This study | - | This study |
| Model resolution (Å) | - | 3.0 (masked) | - | 3.2 (masked) |
| FSC threshold | - | 0.5 | - | 0.5 |
| Model composition | | | | |
| Nonhydrogen atoms | - | 29,557 | - | 29,557 |
| Protein residues | - | 3,788 | - | 3,788 |
| Ligands | - | 2 ATP, ADP, Pi, 2 MG | - | 2 ATP, ADP, Pi, 2 MG |
| Bond length (Å) | - | 0.003 | - | 0.004 |
| Bond Angles (°) | - | 0.547 | - | 0.630 |
| **Validation** | | | | |
| MolProbity score | - | 1.12 | - | 1.17 |
| EMRinger score | - | 3.92 | - | 3.36 |
| Clashscore | - | 3.32 | - | 3.82 |
| Rotamer outlier (%) | - | 0.00 | - | 0.00 |
| CaBALM outlierl (%) | - | 1.36 | - | 1.52 |
| Ramachandran plot | | | | |
| Favored (%) | - | 98.51 | - | 98.59 |
| Allowed (%) | - | 1.49 | - | 1.41 |
| Disallowed (%) | - | 0.00 | - | 0.00 |

**Supplementary Table 5-2. Cryo-EM data collection, refinement and validation statistics for V/A-ATPase obtained under 6 mM ATP condition.**

| | State2-1 | | State2-2 | |
|---|---|---|---|---|
| | $V_oV_1$ | $V_1EG$ | $V_oV_1$ | $V_1EG$ |
| EMDB ID | 31851 | 31853 | 31852 | 31854 |
| PDB ID | | 7VAN | | 7VAO |
| **Data collection and processing** | | | | |
| Magnification | 81,000x | | | |
| EM & Voltage (kV) | Titan Krios, 300 | | | |
| Total dose (e-/Å$^2$) | 50 | | | |
| Pixel size (Å/pix) | 0.88 | | | |
| Defocus range ($\mu$m) | -0.8 to -2.0 | | | |
| Symmetry imposed | C1 | | | |
| Initial particle # | 2,300,834 | | | |
| Final Particle # | 36,693 | | 12,087 | |
| Map resolution (Å) | 3.3 | 3.0 | 4.0 | 3.4 |
| FSC threshold | 0.143 | | | |
| **Refinement** | | | | |
| Initial model used (PDBID) | - | This study | - | This study |
| Model resolution (Å) | - | 3.0 (masked) | - | 3.4 (masked) |
| FSC threshold | - | 0.5 | - | 0.5 |
| Model composition | | | | |
| Nonhydrogen atoms | - | 29,557 | - | 29,557 |
| Protein residues | - | 3,788 | - | 3,788 |
| Ligands | - | 2 ATP, ADP, Pi, 2 MG | - | 2 ATP, ADP, Pi, 2 MG |
| Bond length (Å) | - | 0.005 | - | 0.004 |
| Bond Angles (° ) | - | 0.571 | - | 0.552 |
| **Validation** | | | | |
| MolProbity score | - | 1.25 | - | 1.30 |
| EMRinger score | - | 3.81 | - | 3.03 |
| Clashscore | - | 3.69 | - | 4.48 |
| Rotamer outlier (%) | - | 0.00 | - | 0.00 |
| CaBALM outlierl (%) | | 1.44 | - | 1.47 |
| Ramachandran plot | | | | |
| Favored (%) | - | 97.53 | - | 97.66 |
| Allowed (%) | - | 2.47 | - | 2.34 |
| Disallowed (%) | - | 0.00 | - | 0.00 |

**Supplementary Table 5-3. Cryo-EM data collection, refinement and validation statistics for V/A-ATPase obtained under 6 mM ATP condition.**

| | State3-1 | | State3-2 | |
|---|---|---|---|---|
| | $V_oV_1$ | $V_1EG$ | $V_oV_1$ | $V_1EG$ |
| EMDB ID | 31855 | 31857 | 31856 | 31858 |
| PDB ID | | 7VAP | | 7VAQ |
| **Data collection and processing** | | | | |
| Magnification | 81,000x | | | |
| EM & Voltage (kV) | Titan Krios, 300 | | | |
| Total dose (e-/Å$^2$) | 50 | | | |
| Pixel size (Å/pix) | 0.88 | | | |
| Defocus range (μm) | -0.8 to -2.0 | | | |
| Symmetry imposed | C1 | | | |
| Initial particle # | 2,300,834 | | | |
| Final Particle # | 31,631 | | 5,300 | |
| Map resolution (Å) | 3.3 | 3.0 | 4.5 | 3.6 |
| FSC threshold | 0.143 | | | |
| **Refinement** | | | | |
| Initial model used (PDBID) | - | This study | - | This study |
| Model resolution (Å) | - | 3.1 (masked) | - | 3.6 (masked) |
| FSC threshold | - | 0.5 | - | 0.5 |
| Model composition | | | | |
| Nonhydrogen atoms | - | 29,557 | - | 29,557 |
| Protein residues | - | 3,788 | - | 3,788 |
| Ligands | - | 2 ATP, ADP, Pi, 2 MG | - | 2 ATP, ADP, Pi, 2 MG |
| Bond length (Å) | - | 0.004 | - | 0.003 |
| Bond Angles (°) | - | 0.526 | - | 0.0524 |
| **Validation** | | | | |
| MolProbity score | - | 1.18 | - | 1.29 |
| EMRinger score | - | 3.49 | - | 2.51 |
| Clashscore | - | 3.86 | - | 5.44 |
| Rotamer outlier (%) | - | 0.00 | - | 0.00 |
| CaBALM outlierl (%) | - | 1.66 | - | 1.71 |
| Ramachandran plot | | | | |
| Favored (%) | - | 97.95 | - | 98.19 |
| Allowed (%) | - | 2.05 | - | 1.81 |
| Disallowed (%) | - | 0.00 | - | 0.00 |

**Supplementary Table 6-1. Cryo-EM data collection, refinement and validation statistics for V/A-ATPase obtained under 50 µM ATP condition.**

| | State1 | | |
|---|---|---|---|
| | $V_oV_1$ | $V_1$EG state1-1 | $V_1$EG state1-2 |
| EMDB ID | 31859 | 31860 | 31861 |
| PDB ID | | 7VAR | 7VAS |
| **Data collection and processing** | | | |
| Magnification | 81,000x | | |
| EM & Voltage (kV) | Titan Krios, 300 | | |
| Total dose (e-/Å$^2$) | 50 | | |
| Pixel size (Å/pix) | 0.88 | | |
| Defocus range (μm) | -0.8 to -2.0 | | |
| Symmetry imposed | C1 | | |
| Initial particle # | 1,671,397 | | |
| Final Particle # | 84,150 | 42,167 | 35,872 |
| Map resolution (Å) | 2.7 | 2.9 | 3.0 |
| FSC threshold | 0.143 | | |
| **Refinement** | | | |
| Initial model used (PDBID) | - | This study | This study |
| Model resolution (Å) | - | 3.0 (masked) | 3.1 (masked) |
| FSC threshold | - | 0.5 | 0.5 |
| Model composition | | | |
| Nonhydrogen atoms | - | 29,529 | 29,517 |
| Protein residues | - | 3,787 | 3,787 |
| Ligands | - | ATP, ADP 2 MG | ATP, ADP, 2 MG |
| Bond length (Å) | - | 0.003 | 0.003 |
| Bond Angles (°) | - | 0.536 | 0.502 |
| **Validation** | | | |
| MolProbity score | - | 1.08 | 1.19 |
| EMRinger score | - | 3.92 | 3.52 |
| Clashscore | - | 2.93 | 4.10 |
| Rotamer outlier (%) | - | 0.06 | 0.13 |
| CaBALM outlierl (%) | - | 1.45 | 1.74 |
| Ramachandran plot | | | |
| Favored (%) | - | 98.83 | 98.14 |
| Allowed (%) | - | 1.17 | 1.86 |
| Disallowed (%) | - | 0.00 | 0.00 |

**Supplementary Table 6-2. Cryo-EM data collection, refinement and validation statistics for V/A-ATPase obtained under 50 µM ATP condition.**

| | State2 | | | State3 | |
|---|---|---|---|---|---|
| | $V_oV_1$ | $V_1$EG state2-1 | $V_1$EG state2-2 | $V_oV_1$ | $V_1$EG |
| EMDB ID | 31862 | 31863 | 31864 | 31865 | 31866 |
| PDB ID | | 7VAT | 7VAU | | 7VAV |
| **Data collection and processing** | | | | | |
| Magnification | 81,000x | | | | |
| EM & Voltage (kV) | Titan Krios, 300 | | | | |
| Total dose (e-/Å$^2$) | 50 | | | | |
| Pixel size (Å/pix) | 0.88 | | | | |
| Defocus range (μm) | -0.8 to -2.0 | | | | |
| Symmetry imposed | C1 | | | | |
| Initial particle # | 1,671,397 | | | | |
| Final Particle # | 48,316 | 28,883 | 11,908 | 26,525 | |
| Map resolution (Å) | 3.3 | 3.2 | 3.3 | 3.6 | 2.8 |
| FSC threshold | 0.143 | | | | |
| **Refinement** | | | | | |
| Initial model used (PDBID) | - | This study | This Study | - | This study |
| Model resolution (Å) | - | 3.1 (masked) | 3.5 (masked) | - | 3.4 (masked) |
| FSC threshold | - | 0.5 | 0.5 | - | 0.5 |
| Model composition | | | | | |
| Nonhydrogen atoms | - | 29,529 | 29,517 | - | 29,529 |
| Protein residues | - | 3,787 | 3,787 | - | 3,787 |
| Ligands | - | ATP, ADP, 2 MG | ATP, ADP, 2 MG | - | ATP, ADP, 2 MG |
| Bond length (Å) | - | 0.003 | 0.007 | - | 0.002 |
| Bond Angles (°) | - | 0.507 | 0.670 | - | 0.470 |
| **Validation** | | | | | |
| MolProbity score | - | 1.16 | 1.39 | - | 1.18 |
| EMRinger score | - | 3.36 | 3.02 | - | 3.20 |
| Clashscore | - | 3.69 | 5.14 | - | 3.96 |
| Rotamer outlier (%) | - | 0.00 | 0.00 | - | 0.00 |
| CaBALM outlierl (%) | - | 1.69 | 1.79 | - | 1.47 |
| Ramachandran plot | | | | | |
| Favored (%) | - | 98.19 | 97.39 | - | 98.54 |
| Allowed (%) | - | 1.81 | 2.61 | - | 1.46 |
| Disallowed (%) | - | 0.00 | 0.00 | - | 0.00 |

**Supplementary Table 7-1. Cryo-EM data collection, refinement and validation statistics for V/A-ATPase obtained under 4 mM ATPγS condition.**

| | State1 | | |
|---|---|---|---|
| | $V_oV_1$ | State1-1 $V_1EG$ | State1-2 $V_1EG$ |
| EMDB ID | 31867 | 31868 | 31869 |
| PDB ID | | 7VAW | 7VAX |
| **Data collection and processing** | | | |
| Magnification | 60,000x | | |
| EM & Voltage (kV) | CRYOARM300, 300 | | |
| Total dose (e-/Å$^2$) | 50 | | |
| Pixel size (Å/pix) | 0.81 | | |
| Defocus range ($\mu$m) | -0.8 to -2.0 | | |
| Symmetry imposed | C1 | | |
| Initial particle # | 4,677,284 | | |
| Final Particle # | 140,014 | 45,066 | 49,606 |
| Map resolution (Å) | 2.7 | 2.7 | 2.9 |
| FSC threshold | 0.143 | | |
| **Refinement** | | | |
| Initial model used (PDBID) | - | This study | This study |
| Model resolution (Å) | - | 2.9 (masked) | 3.0 (masked) |
| FSC threshold | - | 0.5 | 0.5 |
| Model composition | | | |
| Nonhydrogen atoms | - | 29,557 | 29,557 |
| Protein residues | - | 3,788 | 3,788 |
| Ligands | - | 2 ATPγS, ADP, 2 MG | 2 ATPγS, ADP, 2 MG |
| Bond length (Å) | - | 0.004 | 0.002 |
| Bond Angles (°) | - | 0.566 | 0.492 |
| **Validation** | | | |
| MolProbity score | - | 1.23 | 1.27 |
| EMRinger score | - | 3.97 | 3.26 |
| Clashscore | - | 2.93 | 5.00 |
| Rotamer outlier (%) | - | 0.03 | 0.03 |
| CaBALM outlierl (%) | - | 1.74 | 1.55 |
| Ramachandran plot | | | |
| Favored (%) | - | 98.57 | 97.98 |
| Allowed (%) | - | 1.43 | 2.02 |
| Disallowed (%) | - | 0.00 | 0.00 |

**Supplementary Table 7-2. Cryo-EM data collection, refinement and validation statistics for V/A-ATPase obtained under 4 mM ATPγS condition.**

| | State2 | | State3 | |
|---|---|---|---|---|
| | $V_oV_1$ | $V_1EG$ | $V_oV_1$ | $V_1EG$ |
| EMDB ID | 31870 | 31871 | 31872 | 31873 |
| PDB ID | | 7VAY | | 7VB0 |
| **Data collection and processing** | | | | |
| Magnification | 60,000x | | | |
| EM & Voltage (kV) | CRYOARM300, 300 | | | |
| Total dose (e-/Å$^2$) | 50 | | | |
| Pixel size (Å/pix) | 0.81 | | | |
| Defocus range ($\mu$m) | -0.8 to -2.0 | | | |
| Symmetry imposed | C1 | | | |
| Initial particle # | 4,677,284 | | | |
| Final Particle # | 35,300 | | 22,646 | |
| Map resolution (Å) | 3.3 | 3.3 | 3.6 | 3.6 |
| FSC threshold | 0.143 | | | |
| **Refinement** | | | | |
| Initial model used (PDBID) | - | This study | - | This Study |
| Model resolution (Å) | - | 3.3 (masked) | - | 3.5 (masked) |
| FSC threshold | - | 0.5 | - | 0.5 |
| Model composition | | | | |
| Nonhydrogen atoms | - | 29,557 | - | 29,557 |
| Protein residues | - | 3,788 | - | 3,788 |
| Ligands | - | 2 ATP$\gamma$S, ADP, 2 MG | - | 2 ATP$\gamma$S, ADP, 2 MG |
| Bond length (Å) | - | 0.004 | - | 0.003 |
| Bond Angles (°) | - | 0.572 | - | 0.534 |
| **Validation** | | | | |
| MolProbity score | - | 1.45 | - | 1.34 |
| EMRinger score | - | 2.87 | - | 2.86 |
| Clashscore | - | 6.16 | - | 6.11 |
| Rotamer outlier (%) | - | 0.00 | - | 0.00 |
| CaBALM outlierl (%) | - | 1.82 | - | 1.76 |
| Ramachandran plot | | | | |
| Favored (%) | - | 97.42 | - | 98.09 |
| Allowed (%) | - | 2.58 | - | 1.91 |
| Disallowed (%) | - | 0.00 | - | 0.00 |

**Supplementary Video 1: Structural transition from state1 of $V_{nucfree}$ to state2 of $V_{2nuc}$.**

The state1 of $V_{nucfree}$ is thermally fluctuating between state1-1 and state1-2. This thermal fluctuation also occurs in $V_{3nuc}$ and $V_{2nuc}$. The $V_{nucfree}$ in ground state is activated by the binding of ATP to the catalytic sites. In $V_{2nuc}$ awaiting ATP binding, binding of ATP onto $AB_{open}$ produces state1 of $V_{3nuc}$. In $V_{3nuc}$, the catalytic events in three AB dimers occurs simultaneously with 120° step of DF shaft, resulting in structural transition of state1of $V_{3nuc}$ to state2 of $V_{2nuc}$.

**Supplementary Video 2: Rotation of $V_1$ moiety driven by ATP hydrolysis.** The state1 of $V_{2nuc}$ is thermally fluctuating between state1-1 and state1-2. The binding of ATP onto $AB_{open}$ produces state1 of $V_{3nuc}$.The catalytic events in three AB dimers in $V_{3nuc}$ occurs simultaneously with 120° step of DF shaft.
